# Carbon Dot-based Fluorescent Antibody Nanoprobes as Brain Tumour Glioblastoma Diagnostics

**DOI:** 10.1101/2021.11.29.470408

**Authors:** Mattia Ghirardello, Radhe Shyam, Xia Liu, Teodoro Garcia-Millan, Imke Sittel, F. Javier Ramos-Soriano, Kathreena Kurian, M. Carmen Galan

## Abstract

The development of efficient and sensitive tools for the detection of brain cancer in patients is of the utmost importance particularly because many of these tumours go undiagnosed until the disease has advanced and when treatment is less effective. Current strategies employ antibodies (Abs) to detect Glial Fibrillary Acid Protein (GFAP) in tissue samples, since GFAP is unique to the brain and not present in normal peripheral blood, and it relies on fluorescent reporters.

Herein we describe a low cost, practical and general method for the labelling of proteins and antibodies with fluorescent carbon dots (CD) to generate diagnostic probes that are robust, photostable and applicable to the clinical setting. The two-step protocol relies on the conjugation of a dibenzocyclooctyne (DBCO)-functionalised CD with azide functionalised proteins by combining amide conjugation and strain promoted alkyne-azide cycloaddition (SPAAC) ligation chemistry. The new class of Ab-CD conjugates developed using this strategy was successfully used for the immunohistochemical staining of human brain tissues of patients with glioblastoma (GBM) validating the approach. Overall, these novel fluorescent probes offer a promising and versatile strategy in terms of costs, photostability and applicability which can be extended to other Abs and protein systems.

## 1. Introduction

Overall less than 20% of brain tumour patients are alive 5 years after diagnosis, in part because they present late with large inoperable tumours.^[1]^ There is an urgent need to develop new sensitive tests of brain tumours to help general practitioners in primary care.^[2]^ The most common malignant primary brain tumour called glioblastoma is characterised by abnormal blood vessels resulting in a leaky Blood Brain Barrier (BBB).^[3]^ Glial Fibrillary Acid Protein (GFAP) is unique to the brain and not present in normal peripheral blood. Antibodies targeting GFAP are used to diagnose gliomas in tissue samples. There is evidence that GFAP crosses the leaky BBB and is an early non-specific peripheral blood biomarker which predates the clinical diagnosis of glioblastoma.^[4]^ However, GFAP levels are too low for routine detection by commonly used protein diagnostic tests such as ELISA, and more sensitive methods for its identification are needed.^[5]^

Fluorescent labelling of proteins is a common strategy to investigate their role and function in cells, tissues and organisms.^[6]^ Traditional efforts rely on the use of molecular dyes, which are usually expensive and predisposed to photobleaching. Alternatively, fluorescent nanoparticles can be tuned to exhibit high stability, sensitivity and specificity for their desired target without the limitations of organic fluorophores and as a result these nanoprobes have found many applications as more robust tools in the areas of bioimaging, drug delivery and diagnostics.^[7]^

Among the different types of luminescent nanomaterials, carbon dots have emerged as a new class of carbon-based fluorescent nanomaterials with semi-spherical morphology, unique optical and physico-chemical properties such as chemical inertness, high water solubility, resistance to photobleaching, low cost of fabrication, and very low cytotoxicity.^[8]^ These carbon based fluorescent nanomaterials have been hailed as alternative probes to semiconductor quantum dots which have been linked to heavy metal toxicity which restricted their use in vivo applications.^[9]^ As a results, the use of carbon-based nanomaterials in biology as a platform for gene delivery,^[10]^ cell imaging,^[11]^ diagnosis,^[12]^ and as theranostics^[13]^ has raised a lot of interest.

CDs can be easily produced via the thermal degradation of readily available substances such as citric acid and ethylenediamine furnishing CDs with high fluorescent quantum yield. The fluorescent excitation and emission pattern of CDs can be tuned by changing the synthetic conditions, variables such as the kind of solvent, temperature and the ratio of precursors used during the preparation may provide different nanoparticles able to emit in different spectral regions.^[8b]^ Therefore, the combination of these features makes of CDs excellent candidates for their use as fluorophores for Abs labelling. However, despite the great advantages offered by CDs for many biosensing applications,^[14]^ their use as fluorophores for Abs labelling is still underdeveloped and Ab conjugation strategies to CDs have not been fully developed for direct clinical applications.^[15]^

To this extent, we envisioned the use of fluorescent carbon dots (CDs) for the development of cheap and photostable probes for Abs functionalization that can be used for the detection of GFAP in clinical samples.

Herein, we describe the development of a practical, low cost and general strategy for the labelling of Abs with fluorescent CDs. This first generation of Ab-CD conjugates combines EDC and strain promoted alkyne-azide cycloaddition (SPAAC) ligation chemistry to generate a new class of Ab-CD conjugates which are robust and photostable (Figure 1). Moreover, the clinical versatility of the novel Ab-probes is demonstrated in the immunohistochemical staining of human brain tissues of patients with glioblastoma GBM.

**Figure 1.**
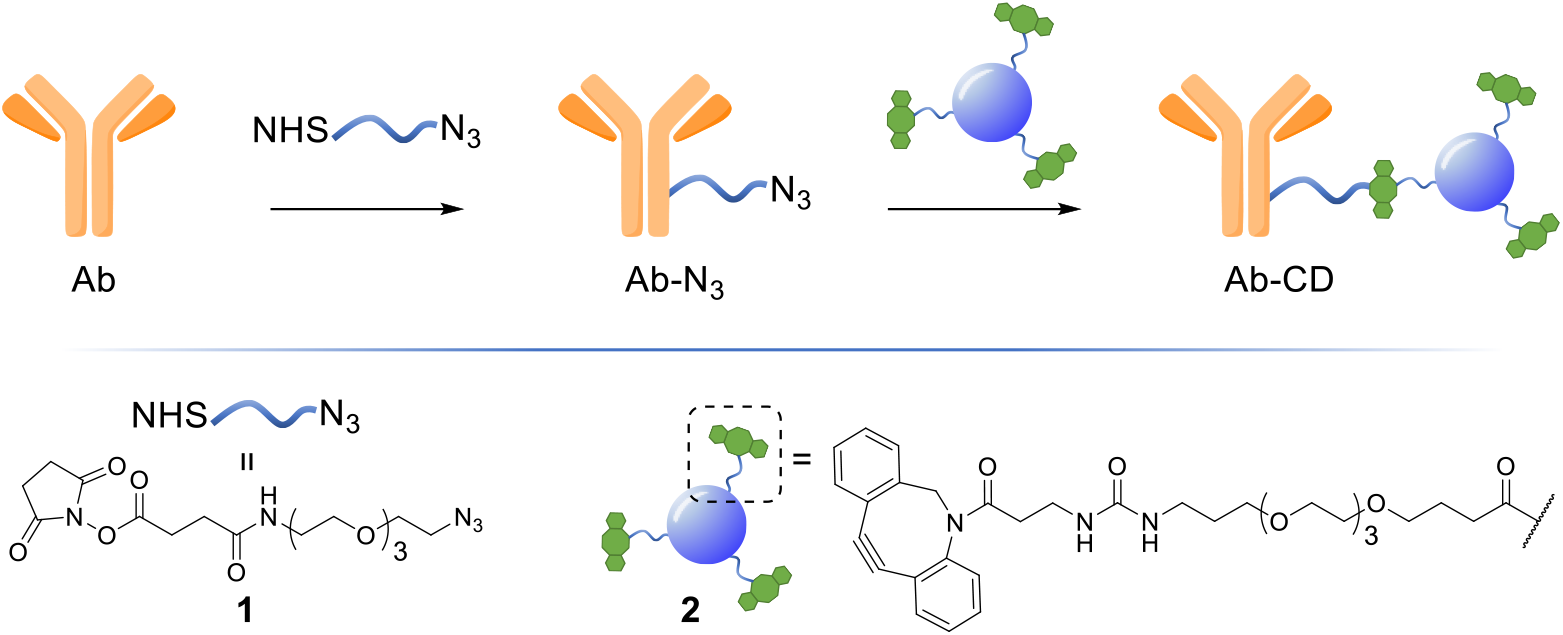
General Ab-CD conjugation strategy.

## 2. Results and Discussion

### 2.1. Synthesis of dibenzocyclooctyne (DBCO)-functionalised fluorescent carbon dots

The synthesis of dibenzocyclooctyne (DBCO)-functionalised CD **2** ready to be conjugated to azido-decorated Abs started from acid functionalised CD **3**, which were prepared in one pot from citric acid and ethylenediamine under microwave irradiation (domestic microwave oven, 300 W) following a modified procedure by Mondal et al.^[16]^ (Figure 2A). The reaction mixture was dissolved in distilled H_2_O and precipitated in an excess of acetone several times to give acid-functionalized CDs, which after dialysis and centrifugal filtration (10 kDa molecular weight cut-off membrane) afforded monodisperse blue emitting nanoparticles as evidenced by fluorescence measurements (Figure 2B). TEM revealed the presence of quasi-spherical nanoparticles with an average size between 2 - 5 nm (N = 262) and a lattice interspacing of 0.34 nm (Figure 2C) which correlate to a graphite core structure.^[17]^

**Figure 2.**
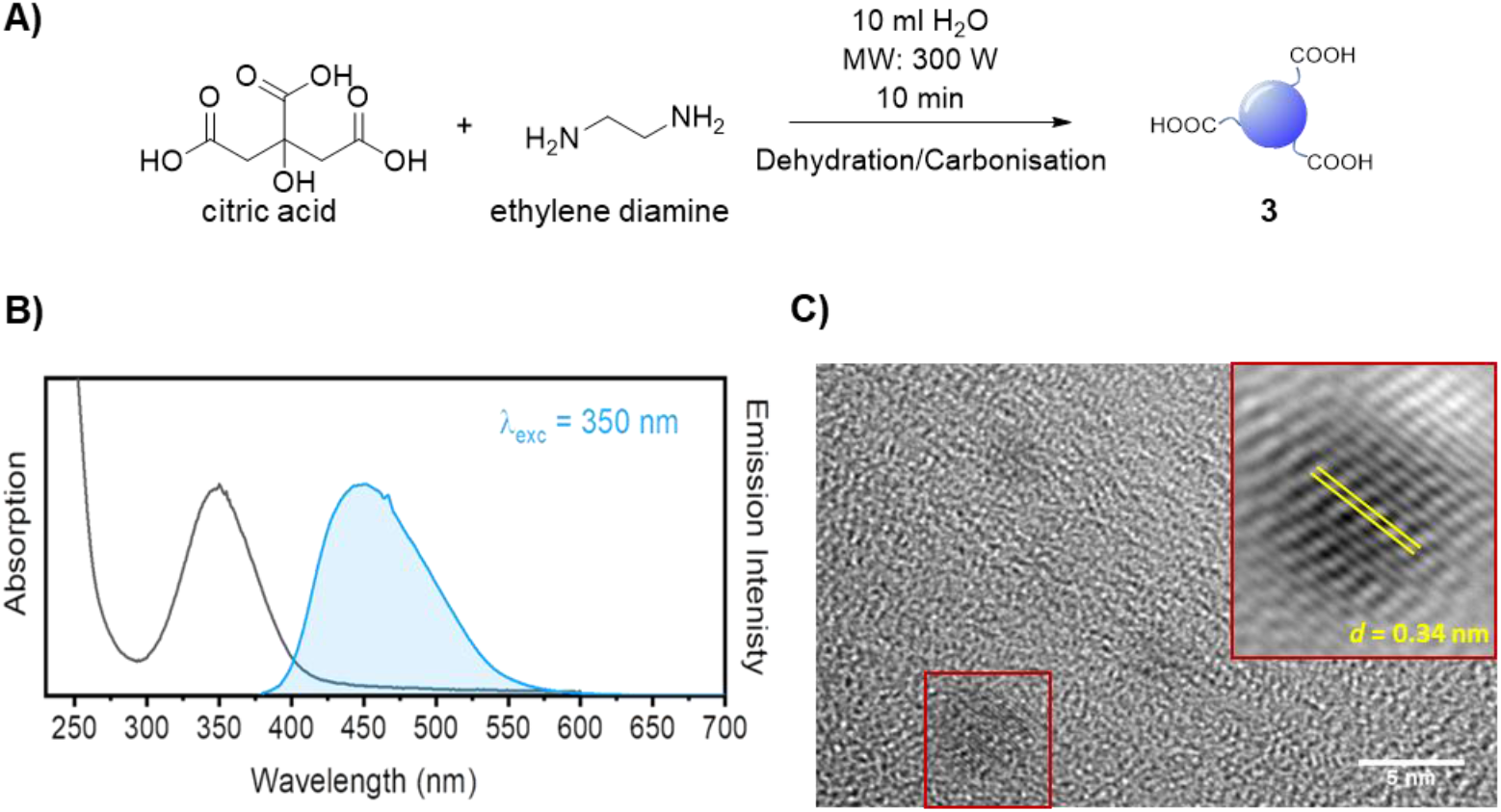
A) Synthetic approach for the synthesis of acidic coated CDs **3**. B) Absorption and emission spectra of CD **3**. C) TEM image of CD **3**. Lattice interspacing (*d*) for single dots is included.

**Figure 2.**
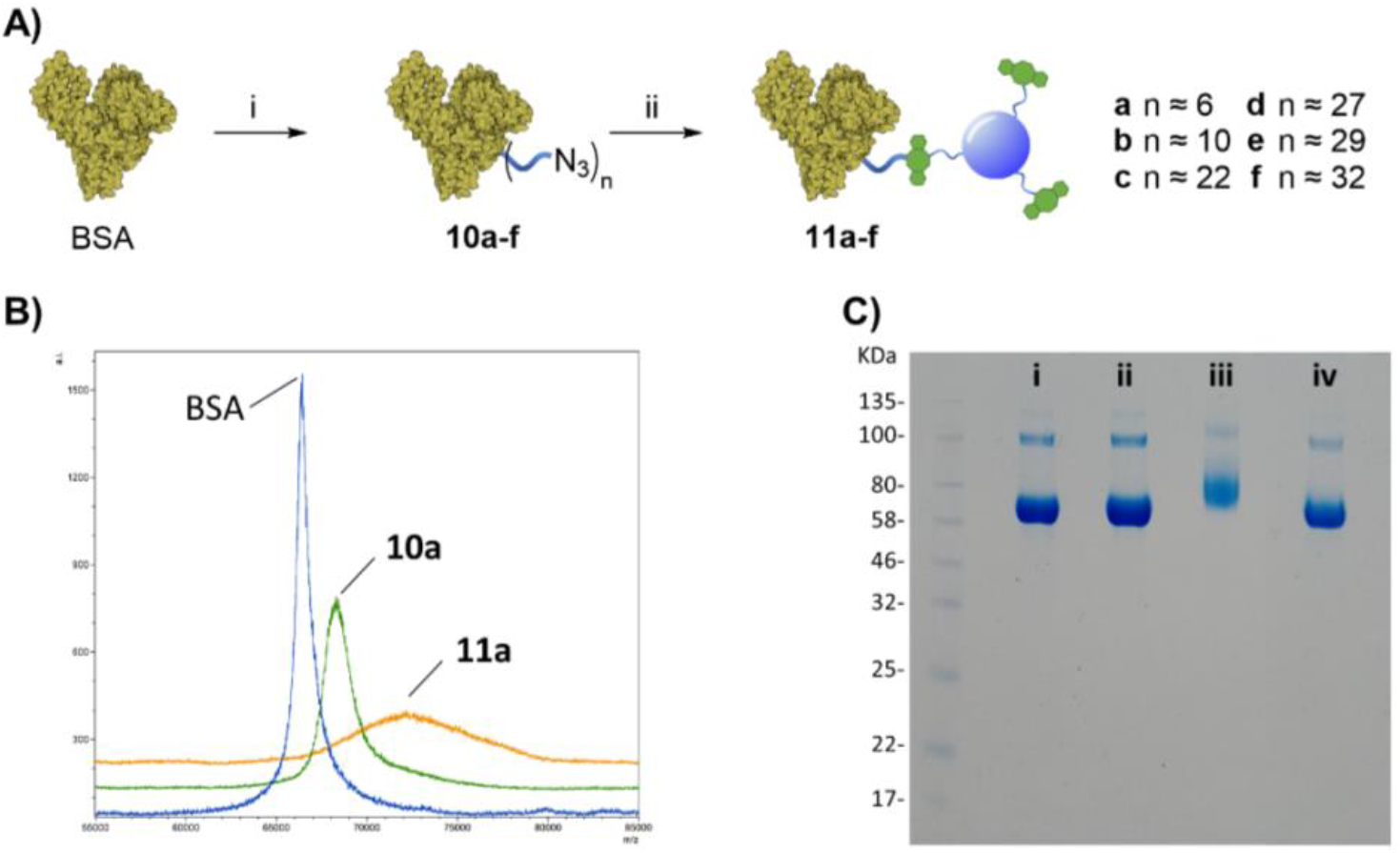
A) *Reagents and conditions:* i) **1**, PBS, 4 h, rt; ii) **2**, 16 h, rt. B) MALDI spectra of BSA, **15a**, and **16a**. Intensity is expressed as arbitrary units (a.i.). C) NuPAGE Gel electrophoresis of BSA derivatives: i) native BSA, ii) **10a**, iii) **11a**, and iv) native BSA (no N_3_ present) mixed with **2**.

Functionalization of CD **3** with DBCO-linker **9**, which was prepared in 4 steps and 47% overall yield, afforded DBCO-CDs **2** (Scheme 1). In brief, mono-amine protection of commercially available 4,7,10- trioxa-1,13-tridecanediamine **4** with Boc_2_O in CH_2_Cl^2^ gave **5**^[18]^ in 99% yield. The free amine in **5** was then reacted with 4-nitrophenyl chloroformate to form activated carbamate **6**, which could then be treated with commercial DBCO-amine **7** to give **8** in 55% yield over the 2 steps. Boc deprotection in the presence of TFA/CH_2_Cl_2_ afforded **9** ready for CD conjugation. HATU mediated nanoparticle functionalization of acid coated CD **3** with **9** was carried out using a 1:0.5 w/w ratio of **3**:**9**, which was found to be optimum to ensure the nanoparticles remained in solution despite the hydrophobic coating. Furthermore, ^1^H, HSQC and Diffusion Ordered (DOSY) NMR spectroscopy analysis demonstrated the successful conjugation of DBCO moieties on the CD (See ESI Figures S11-13).

**Scheme 1.**
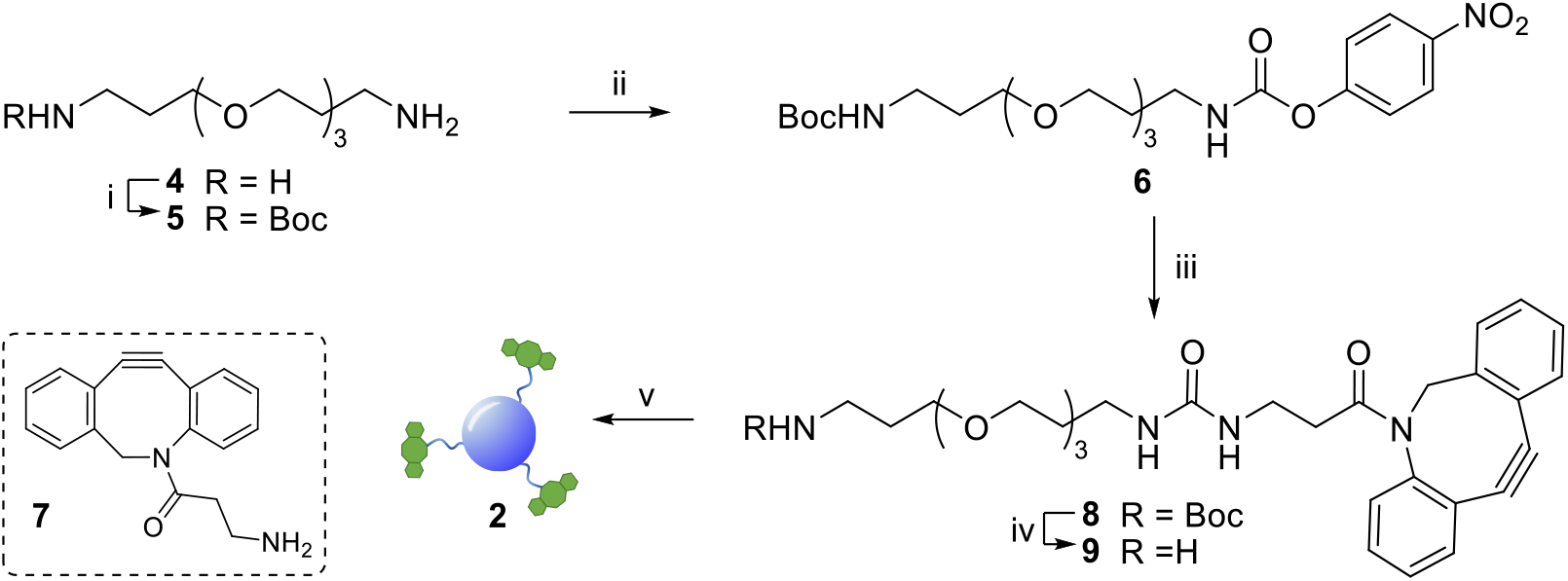
Reagents and conditions: i) Boc_2_O, DCM, 4 h, 0 °C to rt, 99%; ii) 4-nitrophenyl chloroformate, Py, DCM, 3 h, 0 °C to rt, 87%; iii) **7**, Py, DIPEA, DMF, 63%; iv) TFA, DCM, 1.5 h, rt, 86%; v) **3**, HATU, DIPEA, DMF, 5 h, rt.

### 2.2. Protein Conjugation Strategy

There are several antibody-drug conjugation strategies available which include: amide formation, periodate oxidation of carbohydrates in the Fc region and subsequent functionalization of the novel aldehyde function produced within the glycosidic moiety, click reactions involving 1,3-dipolar cycloadditions, thiol maleimide conjugations and thiol-mediated alkylation among others.^[19]^ Among those, the EDC/NHS-promoted amide formation by targeting primary amines found on lysine residues, appears to be one of the most common and direct approaches for nanoparticle loading. ^[14, 20]^ Control over the Ab/nanoparticle loading ratio to avoid the over functionalization of the Ab, as well as the type and length of spacer molecule between the probe and Ab are essential to avoid loss of protein’s pharmacokinetic properties. Moreover, excessive labelling may also interfere with the antigen-biding event. In this context, an Ab/drug ratio of around 1:4 is found to be optimal^[21]^ and this parameter was considered in the preparation of our Ab-CD conjugates. NHS activated conjugation represents the golden standard as it allows for a fast and reliable Abs covalent functionalization, NHS-functionalised molecules react with solvent exposed primary amines such as lysine residues on the surface of the Abs.^[22]^ Taking into account the above considerations, NHS-linker **1** (for its preparation see ESI section 3.3)^[23]^ featuring an azido motif that can selectively react with the DBCO-CD **2**. Moreover, both **1** and **2** were designed with a triethylene glycol spacer to help with water solubility and to reduce steric hindrance between the Ab and the surface of the CD.

#### 2.2.1 BSA-CD conjugation strategy

With both CD-DBCO **2** and NHS-linker **1** in hand, the feasibility of our labelling approach was initially evaluated on bovine serum albumin (BSA) protein, as an inexpensive model system (Figure 2A). In the first step, the protein was functionalised with the azido-containing linker **1**, in brief a solution of BSA in PBS (100μL, 36.1 μM) was reacted with excess amounts of **1** (Figure 2A, different molar excess of **1** were tested, entries a-f: from 40 to 1333 eq.) under gentle shaking at room temperature for 4 h. Removal of the excess of **1** and washes via spin filtration over 30 KDa cut-off membrane gave **10a-f** with different degrees of functionalization from approximately 6 to a maximum of 32 linker units per protein as determined by Matrix Assisted Laser Desorption/Ionization (MALDI) mass spectrometry using the spotting procedure described by Signor et al.^[24]^ (See figure 2A, and ESI: figure S20 and Table S1).

Once the protein is decorated with the azido functionalities, chemoselective Cu-free click conjugation with DBCO-CD **2** was attempted in PBS by mixing **10a** (BSA with 6 N_3_-linker units) and **2** at room temperature for 16 h. A 5-fold excess of **2** in weight with respect to the protein was used to ensure all the available N_3_ moieties were conjugated. Following spin filtration (30 KDa cut-off membrane) to remove the excess of **2**, BSA-CD conjugate **11a** was obtained as determined by MALDI (Figure 2B). The conjugation of CDs to BSA caused a shift toward larger molecular weights and a broadening of the peak as expected from protein conjugation with a disperse nanoparticle system such as our CDs.^[25]^

Gel electrophoresis was also used to further confirm the effective BSA-CDs conjugation, by allowing us to compare the MW of the different protein adducts (Figure 2C). Whereas the addition of 6 low molecular weight (MW) N_3_-linkers on **10a** did not show any significant changes on the gel when compared to with native BSA (Figure 2C, i *vs* ii), a noticeable increase in MW was shown for the BSA- CD adduct **11a** (Figure 2C, i *vs* iii), which further validates the MALDI data. Gel electrophoresis analysis of BSA-N_3_ derivatives **10b-f** was also possible (see ESI figure S22).

Moreover, to exclude the possibility of non-specific BSA adsorption on CD nanoparticles, unfunctionalized BSA, which lacks azido motifs, and DBCO-CDs were pre-mixed together and run on the same well showing no MW changes with respect to BSA alone (Figure 2C, i *vs* iv) which demonstrated the chemospecific labelling of the protein via SPAAC reaction.

#### 2.2.2 Anti-GFAP Abs-CD conjugation

Having demonstrated successful protein labelling with our strategy, CD conjugation on clinically relevant rabbit polyclonal anti-glial fibrillary acidic protein antibodies (anti-GFAP Abs) was next attempted. NHS-linker **1** conjugation to anti-GFAP Abs was performed as before (Figure 3A). A solution of Abs in PBS (11.1 μM) was treated with an excess of **1** (Figure 3A, entries a-d: 0.12 – 2.41 μmol) and left shaking at room temperature over 4 h. MALDI analysis of the products showed peak broadening and shifts towards higher MWs for the azido functionalised anti GFAP Abs which could be used to estimate the average degree of substitution (Figure 3B, and ESI: Figure S24 and Table S2). In general, it was found that a maximum of 30 azido containing linkers could be conjugated to the Abs at the higher concentrations, while a degree of functionalization of 4 linker moieties was achieved when 100 molar equivalents of **1** was used for the conjugation, which is optimum to maintaining Ab function and good pharmacokinetic and toxicology profile.^[21]^ The anti-GFAP Abs-N_3_ derivatives **12a-d** were then treated with an excess of DBCO-CD **2** as previously described for BSA and following spin filtration over 50 KDa cut-off membrane to remove the excess of unconjugated **2**, and anti-GFAP Abs-CD conjugates **13a-d** were generated. Gel electrophoresis was used to confirm CD labelling of the anti GFAP-Abs. As for the BSA model gel electrophoresis showed negligible differences in terms of MW for **12a-d** when compared to native Abs (Figure S25 i *vs* iii-vi). It is worth noting that although MALDI clearly shows MS differences between native and azido functionalised Abs, no changes on MW were detectable on the gels for neither anti-GFAP Abs or anti-GFAP Abs-N_3_ since the molecular weight differences between the species is negligible at the level of MW resolution for gel electrophoresis. Indeed, the different Abs-CD probes showed significant increase in MW for **13a-d**, which correlated to their degree of azido functionalization, when compared to native anti-GFAP Abs (Figure 3C).^[26]^ To confirm that the CD labelling of the Abs is not due to non-specific interactions, fluorescence images of a native Abs and Abs-N_3_ that were treated with DBCO-CD **2** prior to purification showed fluorescent labelling only for azido containing Abs as expected (Figure 3D). Moreover, as previously demonstrated for BSA, gel electrophoresis of unfunctionalized Abs were pre-mixed with **2** and run on the same well showed no MW changes with respect to Abs alone (Figure S25, i *vs* ii) confirming the absence of non-specific interactions between the CD and the Abs.

**Figure 3.**
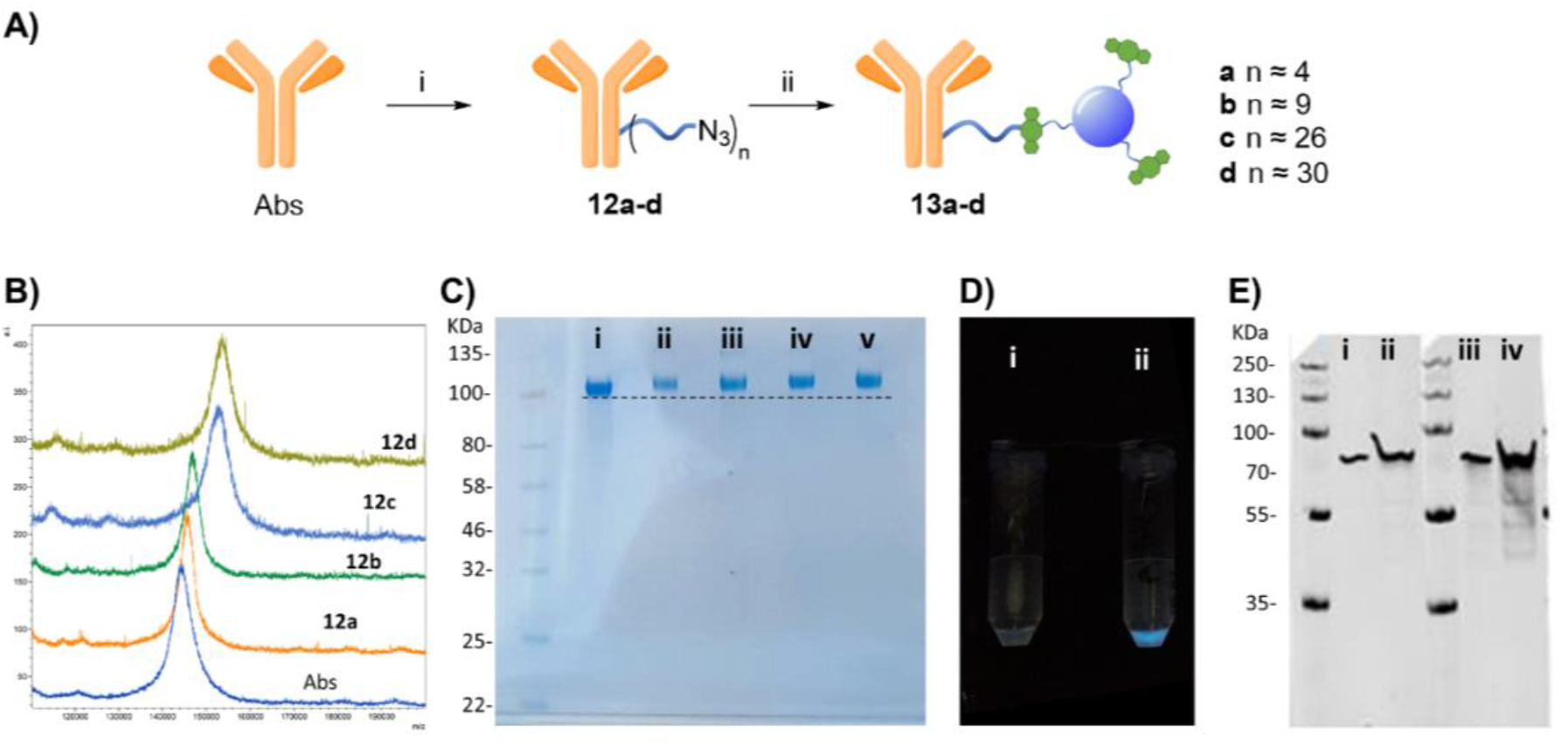
A) *Reagents and conditions:* i) **1**, PBS, 4 h, rt; ii) **2**, 16 h, rt. B) MALDI spectra of Abs, and azide derivatives **17a-d**, Intensity is expressed as arbitrary units (a.i.). C) NuPAGE Gel electrophoresis of Abs derivatives: i) native Abs, ii) **13a**, iii) **13b**, iv) **13c** and v) **13d**. D) Comparison under the UV light of the fluorescent emission of i) non-functionalise Abs mixed with **2** and purified, and ii) purified **13a**. E) Western blot using recombinant human GFAP protein and **13a**, i) 20 ng of GFAP and **13a** 1:1000 dilutions, ii) 100 ng of GFAP and 13a 1:1000 dilutions, iii) 20 ng of GFAP and unlabelled Abs, iv) 100 ng of GFAP and unlabelled Abs, IRDye 680RD goat anti-rabbit IgG was used as secondary antibody.

Furthermore, western blot analysis using human GFAP with **13a** (functionalised with 4 linker units) was used to demonstrate the novel anti-GFAP Ab-CD **13a** adducts retained their ability to recognise the target antigen. To that end, a goat anti-rabbit secondary antibody equipped with a near IR probe was used on the western blot, confirming the presence of the rabbit anti-GFAP Abs bound to the human GFAP antigen (Figure 5E).^[27]^

### 2.3. Anti-GFAP Abs-CD Immunostaining of Clinical Tissue Brain Cancer Patient Samples

GFAP immunostaining is the most commonly used method to examine the distribution of astrocytes and the hypertrophy of astrocytes in response to neural degeneration or injury as in the development of glioblastoma.^[4]^ To demonstrate the versatility of our CD-based Ab labels for diagnosis applications, we have examined GFAP in 13 formalin-fixed paraffin embedded biopsy brain tumour samples from different patients (12 glioblastoma, IDH wildtype, WHO Grade 4 and 1 negative control schwannoma, WHO Grade I, see ESI: table S3) using our conjugated antibody **13a** (Figure 4). We identified immunofluorescence within all the glioblastoma cases (as assessed by a consultant neuropathologist KMK) using the conjugated anti-GFAP antibody **13a** (Figure 4A and 4B, for the complete set of pictures see ESI Figure S27). We identified the correct pattern of cytoplasmic staining (blue) of the GFAP intermediate filament in the glioblastoma cell cytoplasm (Figure 4A and 4B). The intensity and extent of GFAP immunopositivity showed inter and intra-tumoural heterogeneity in keeping with known biological variation between cases. The negative control schwannoma showed no positive staining using the conjugated GFAP antibody as expected (Figure 4C). There was no variation of GFAP staining with age, sex or molecular parameters within the small cohort as expected. In addition, control labelling experiments of glioblastoma samples with CD-DBCO **2** without the Abs, showed no labelling further demonstrating that Abs-CD **13a** is responsible for the labelling observed (see ESI, Figure S26A).

**Figure 4.**
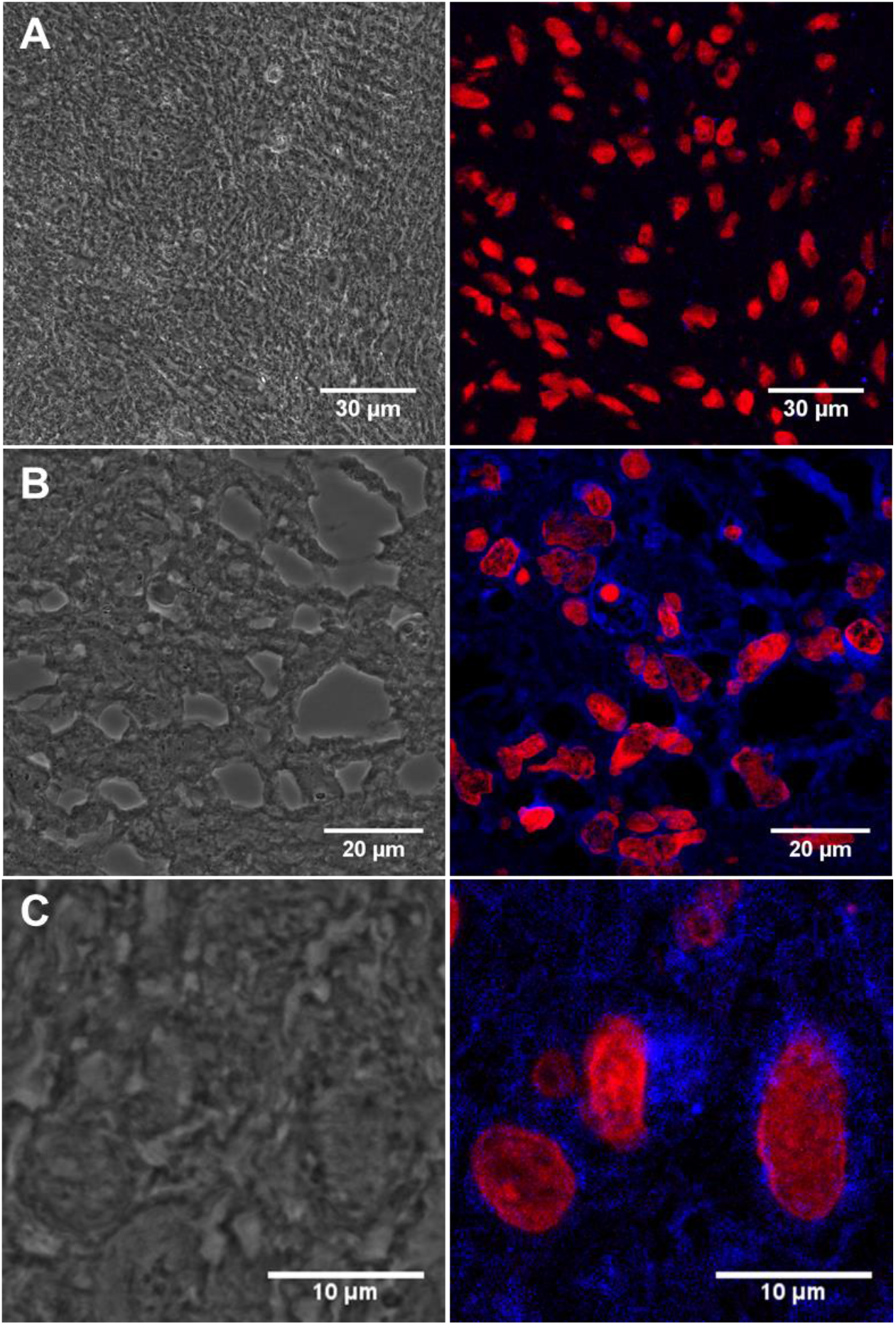
A) Confocal image of the negative control benign schwannoma stained with Anti-GFAP Abs-CD **13a** showing immunofluorescence red labelling nuclei and negative for blue GFAP staining (patient ID: N1214A1). B) Immunostained tissue section of malignant brain tumour glioblastoma stained with **13a** showing immunofluorescence red labelling nuclei and blue labelling cobweb pattern of intermediate filament GFAP (patient ID: 08/0160b). Magnified view of **13a** labelled tissue section of malignant brain tumour glioblastoma showing blue immunopositivity in a single cell (patient id: 10/0053). Left panel = bright field showing monolayer. Right panel = Anti-GAFP Abs-CD probe **13a** labelled tissue. For full scale pictures of A B and C refer to ESI, Figure S27 A, K and G respectively.

## 3. Conclusions

In summary, we have successfully developed a new class of carbon dot-based fluorescence labels that can be “clicked” onto suitably functionalised proteins such as Abs in a chemoselective manner. The two-step strategy relies on the used DBCO-functionalised CDs and alkyne-functionalised proteins that can be easily prepared by simple amide conjugation methods from suitably functionalised linkers with control over degree of functionalization. The novel anti-GFAP Abs-CD probes developed here retained their ability to interact with the human GFAP. Moreover, we have demonstrated our novel probes show reliable binding in a range of clinical malignant brain tumour glioblastoma cases, in tissue sections. Overall, this new class of probes offer a promising and versatile strategy in terms of costs, photostability and applicability which can be extended to other Abs and protein systems. This type of cheap and rapid nanoparticle test have the potential to pave the way for novel strategies to identify the presence of tumour markers such as GFAP in clinical samples to support early diagnosis of brain tumours in primary care. Early diagnosis would potentially improve survival and reduce anxiety in these patients by giving them more surgical and treatment options earlier in the course of their disease.

## 4. Materials and methods

### 4.1 General Experimental

Reagents and solvents were purchased as reagent grade from Sigma Aldrich or ThermoFisher and used without further purification. For column chromatography, silica gel 60 (230-400 mesh, 0.040-0.063 mm) was purchased from Merck and for gel filtration Sephadex G-25 from GE Healthcare. Thin Layer Chromatography (TLC) was performed on aluminium sheets coated with silica gel 60 F254 purchased from Merck. Dialysis was performed with Cole-Parmer Spectra Por Dialysis Tubing, 500-1000 MWCO. Centrifugal spin filtration was performed on Amicon Ultra-0.5 mL purchased from Merck. Carbon dots were prepared using a domestic microwave oven (300 W). NMR spectra were recorded on Bruker AV 400 MHz or AV 500 MHz spectrometers, using the residual solvent peaks as internal reference at 298 K. Chemical shifts are reported as parts per million and coupling constants (J) given in Hertz. All the assignments were confirmed by one- and two-dimensional NMR experiments (DEPT, COSY, HSQC). Mass spectra were obtained by the University of Bristol mass spectrometry service using electrospray ionisation (ESI) acquired on a Micromass LCT mass spectrometer or aVG Quattro mass spectrometer and MALDI spectra were acquired on Bruker ultrafleXtreme 2 (TOF). Zeta potential analysis was carried out using Malvern Instruments Nano-Z ZEN 2600 and conducted in distilled H_2_O at a concentration of 4 mg·mL^-1^. BSA was purchased from ThermoFisher (23209). Polyclonal Rabbit Anti-Glial Fibrillary Acidic Protein was purchased from Agilent Dako (Z033401-2). Recombinant Human GFAP protein, used in the Western Blot test was purchased from abcam (ab114149). Secondary IRDye^®^ 680RD Goat anti-Rabbit IgG used in the Western Blot test were purchased by LI-COR. Human brain tissue samples were kindly provided by the Southmead Hospital, (University of Bristol, UK).

### 4.2 Chemical synthesis

#### Compound 6

To a stirred solution of 4-nitrophenyl chloroformate (2.98 g, 14.79 mmol) and Py (1 mL, 12.33 mmol) in anhydrous DCM (70 mL) at 0 °C, a solution of **5** (1.58 g, 4.93 mmol) in dry DCM (20 mL), was added over 1 h at 0 °C. Once the addition was completed, the solution was stirred for further 2 h at room temperature. The reaction was quenched by the addition of saturated aq. NH_4_Cl solution (50 mL) and the mixture was extracted with DCM (3 x 50 mL). The combined organic phases were dried with anhydrous MgSO_4_, filtered and concentrated under reduced pressure. The residue was purified by column chromatography on silica gel (Hex/EtOAc 1:0 to 3:7, v/v) furnishing **6** (2.08 g, 87% yield) as a transparent syrup. **^1^H NMR** (500 MHz, Chloroform-*d*) δ 8.27 – 8.20 (m, 2H, Ar), 7.35 – 7.27 (m, 2H, Ar), 6.12 (s, 1H, NH), 4.89 (s, 1H, NH), 3.69 – 3.62 (m, 8H, OCH_2_), 3.59 (dd, *J* = 5.8, 3.5 Hz, 2H, OCH_2_), 3.51 (t, *J* = 6.0 Hz, 2H, OCH_2_), 3.42 (q, *J* = 6.0 Hz, 2H, NCH_2_), 3.21 (q, *J* = 6.5 Hz, 2H, NCH_2_), 1.87 (h, *J* = 6.0 Hz, 2H, CH_2_C*H*_2_CH_2_), 1.77 – 1.69 (m, 2H, CH_2_C*H*_2_CH_2_), 1.43 (s, 9H, CH_3_). **^13^C NMR** (126 MHz, CDCl_3_) δ 156.3, 156.2, 153.4, 144.8, 125.2, 122.1, 79.2, 70.7, 70.7, 70.4, 70.3, 70.1, 69.7, 40.2, 38.6, 29.8, 29.1, 28.6. **HRMS (ESI)** m/z: Calcd for C_22_H_35_N_3_O_9_Na (M+Na)^+^ 508.2265, found 508.2279

#### Compound 8

To a stirred solution of **6** (81 mg, 0.29 mmol) and Py (0.5 mL, 6.21 mmol) in anhydrous DCM (5 mL), a solution of DBCO-amine **7** (286 mg, 0.59 mmol) in anhydrous DCM (2 mL) was added dropwise at room temperature, followed by the addition of DIPEA (153 μL, 0.88 mmol) at room temperature. The solution was stirred for 4 h at room temperature, then diluted with DCM (100 mL), washed wit saturated aq. NH_4_Cl solution (2 x 50 mL) and brine (1 x 50 mL). The organic phase was dried with anhydrous MgSO_4_, filtered, and concentrated under reduced pressure. The residue was purified by column chromatography on silica gel (EtOAc/MeOH 1:0 to 9:1, v/v) furnishing **8** (116 mg, 63% yield) as a transparent oil. **^1^H NMR** (500 MHz, Chloroform-*d*) δ 7.66 (d, *J* = 7.6 Hz, 1H, Ar), 7.42 - 7.23 (m, 7H, Ar), 5.12 (d, *J* = 13.9 Hz, 1H, CH_2a_^DBCO^), 5.05 (s, 1H, NH), 5.00 (d, *J* = 6.3 Hz, 1H, NH), 4.86 (s, 1H, NH), 3.69 – 3.47 (m, 13H, CH_2b_^DBCO^, OCH_2_), 3.30 – 3.10 (m, 6H, NCH_2_), 2.56 – 2.48 (m, 1H, COCH_2_), 1.98 – 1.84 (m, 1H, COCH_2_), 1.79 – 1.63 (m, 4H, NCH_2_C*H*_2_CH_2_), 1.42 (s, 9H, CH_3_). **^13^C NMR** (126 MHz, CDCl3) δ 172.5, 158.3, 156.1, 151.2, 148.1, 132.1, 129.2, 128.6, 128.2, 128.2, 127.7, 127.1, 125.5, 123.1, 122.5, 114.7, 107.9, 70.5, 70.4, 70.1, 69.8, 69.5, 69.4, 55.5, 38.4, 36.1, 35.6, 29.6, 29.5, 28.4. **HRMS (ESI)** m/z: Calcd for C_34_H_47_N_4_O_7_Na (M+Na)^+^ 645.3259, found 645.3236.

#### Compound 9

Compound **8** (116 mg, 0.19 mmol) was dissolved in a DCM/TFA solution (7 mL, 95:5, v/v) and stirred for 1.5 h at room temperature. The reaction was concentrated under reduced pressure and the residue was purified by column chromatography on silica gel (CHCl_3_/MeOH 1:0 to 95:5, containing a 0.5% of 35% aq. HN_4_OH solution v/v/v) furnishing **9** (84 mg, 86% yield) as a pale brown oil. **^1^H NMR** (500 MHz, D_2_O^25°C^) δ 7.63 (d, *J* = 7.7 Hz, 1H, Ar), 7.46 – 7.35 (m, 6H, Ar), 7.29 – 7.23 (m, 1H, Ar), 5.02 (d, *J* = 14.5 Hz, 1H, CH_2a_^DBCO^), 3.72 – 3.61 (m, 11H, CH_2b_^DBCO^, OCH_2_), 3.52 (t, *J* = 6.4 Hz, 2H, OCH_2_), 3.14 – 2.95 (m, 6H, NCH_2_), 2.32 – 2.16 (m, 2H, CH_2_CO), 1.94 (dt, *J* = 13.5, 6.4 Hz, 2H, NCH_2_C*H*_2_CH_2_), 1.67 (p, *J* = 6.6 Hz, 2H, NCH_2_C*H*_2_CH_2_). **^13^C NMR** (126 MHz, D_2_O^25°C^) δ 174.3, 159.7, 150.6, 147.7, 131.9, 129.1, 129.1, 128.9, 128.5, 128.1, 127.0, 125.7, 122.4, 121.6, 114.3, 107.8, 69.6, 69.5, 69.4, 69.3, 68.5, 68.3, 55.5, 37.6, 36.8, 36.3, 34.7, 29.0, 26.5. **HRMS (ESI)** m/z: Calcd for C_29_H_39_N_4_O_5_ (M+H)^+^ 523.2915, found 523.2927

#### Carbon Dot 3

Citric acid (1.00 g, 5.2 mmol) was dissolved in distilled H_2_O (10 mL) in a 250 mL conical flask. Ethylenediamine (EDA, 384 μl, 5.72 mmol) was then added to the solution and stirred for 30 min to ensure homogeneity. The conical flask was then placed in a domestic microwave 300 W (inside a fume cupboard) and the solution was reacted for 10 min. A viscous amber residue was obtained which was washed with a solution MeOH:Acetone 1:1 (4xtimes). The residue was then phase-separated by centrifugation and re-dissolved in 15 ml of distilled H_2_O. The CD solution was dialysed in H_2_O using 0.5-1 KDa MWCO Biotech Cellulose Ester membrane. The concentrate CD solution was then lyophilised to yield 1.1 g of CD as an amber powder. To remove high MW components the 100 mg of CD were redissolved in H_2_O and filtered over Amicon Ultra spin filtration (10 KDa cut-off membrane) and liophilized furnishing 92 mg of CD as an amber powder. Procedure modified from the one reported by Mondal et al.^[16]^ See SI for full characterization.

#### DBCO-CDs 2

To a stirred solution of CDs **3** (18.4 mg) in dry DMF (1.84 mL), HATU (13.4 mg, 0.035 mmol) and DIPEA (6.1 μL, 0.035 mmol) were added and the solution was allowed to stir for further 15 minutes at room temperature. A solution of **9** (9.2 mg, 0.018 mmol) in dry DMF (0.5 mL) was added and the solution was stirred at room temperature for 5 h. H_2_O (0.5 mL) was then added to quench the reaction and the solution was stirred for further 10 minutes at room temperature and concentrated under reduced pressure. The residue was redissolved in aq. 0.1 M NaOH solution (3 mL) and stirred for 1 h at room temperature. The pH was neutralized with the addition of aq. HCl 1M solution (0.15 mL), diluted with H_2_O (20 mL), washed with Et_2_O (5 x 10 mL), and the water phase was concentrated under reduced pressure. The residue was purified via 1 KDa cut-off dialysis membrane against water, changing the water bath 3 times over a 24 h period. The purified solution was then freeze-dried furnishing **2** (10.2 mg) as a pale yellow solid.**^1^H NMR** (500 MHz, D_2_O, 25 °C) characteristic resonances δ ppm = 7.77 – 7.06 (m, Ar), 5.14 – 5.04 (m, CH_2_^DBCO^a) 4.29 – 2.50 (PEG linker and CH_2_^DBCO^b). **^1^H /^13^C NMR HSQC** (126 MHz, D_2_O 25 °C) characteristic resonances δ ppm = 131.6 (Ar), 127.1 (Ar), 129.1 (Ar), 125.8 (Ar), 55.5 (CH_2_^DBCO^), 44.5, 41.9, 69.3, 68.4, 39.0, 38.4, 36.1, 36.4, 36.4, 43.8, 44.2, 36.4, 36.4, 34.5, 28.3, 19.6, 0.6

### 4.3. General protein/Abs conjugation procedure

#### Step 1 Azide functionalization

To a solution of protein in PBS (100 μL,36.1 μM for BSA or 100 μL, 11.1 μM Abs), different amounts of compound **1** (0.1 mg/μL in DMSO stock solution) from 0.14 to 4.81 μmol for BSA and 0.12 – 2.41 μmol for Abs, were added, respectively (See Table S1 and Table S2). The final solution was mixed in a shaker at 400 rpm for 4 h at room temperature. The product was purified via spin-filtration using 30 KDa or 50 KDa cut-off membrane for BSA or Abs respectively, at 4000 g per 20 minutes. The concentrated protein solution was diluted with 100 μL of PBS and concentrated again; this washing step was repeated two more times to remove unbound linker **1** and by-products of the reaction, furnishing a concentrated **10a-f** or **12a-d** for BSA and Abs derivatives respectively.

#### Step 2 CD-conjugation

The concentrated **10a-f** or **12a-d** solution prepared in Step 1 was diluted with 100 μL of a PBS solution containing DBCO-CD **2** (2 mg/mL) mixed in a shaker at 400 rpm for 16 h at room temperature. The product was purified via spin-filtration using 30 KDa or 50 KDa cut-off membrane for BSA or the Abs respectively, at 4000 g per 20 minutes. The concentrated protein solution was diluted with 100 μL of PBS and concentrated again; this washing step was repeated three more times to remove the excess of **2** (4 washing steps were judged enough to remove the excess of **2** since no fluorescence was detected by the naked eye in the washing solution passing through the membrane under UV lamp in the last wash), furnishing a concentrated **11a-f** or **13a-d** solution

### 4.4. Gel Electrophoresis

SDS-PAGE on 4-12% NuPage gels (Life Technologies) was performed for labelled and unlabelled protein/Abs samples. Loading dye was added to each sample and heated at 100°C for 5 minutes. SeeBlue™ (ThermoFisher Scientific) was used as a ladder. Lanes were loaded at similar protein concentrations and the gel was run with MES buffer at 180V for 40 minutes. The images were acquired after staining with Pageblue (ThermoFisher Scientific/Pierce) protein staining solution.

### 4.5. Western Blot analysis

Recombinant human GFAP protein were run on 12% SDS-PAGE and transferred onto membrane via Trans-Blot Turbo Transfer System (BIO-RAD). Membrane was incubated in blocking buffer (PBS, 0.1% Tween and 2% milk) for one hour and then incubated in primary anti-GFAP antibodies or anti-GFAP- CD conjugates **13a** (results and discussion) overnight in the cold room. Membranes were washed and then incubated in secondary antibodies for one hour. The membranes were washed and then visualized on a LI-COR Odyssey imaging system.

### 4.6. Tissue staining protocol for GFAP immunofluorescence with Abs-CD 13a

The tissue sections were deparaffinized and rehydrated as follows, the sections were incubated in three washes of xylene for 2 min each, followed by two washes of 100%, 95% ethanol for 10 min each. The sections were then washed twice in distilled H_2_O for 5 min each.

The tissue slides were then placed in the microwaveable vessel. Tris-EDTA antigen retrieval buffer (10 mM Tris base, 1 mM EDTA solution, 0.05% Tween 20, pH 9.0) was added and placed inside a dedicated domestic microwave microwave, which was set to full power (950 W) until the solution came to a boil. The solution was boiled for 20 min from this point and left on the bench at room temp to cool for 30 mins. The slides were then washed 2 x 5 min with TBS plus 0.025% Triton X-100 with gentle agitation. The slides were blocked in Superblock buffer (Thermofisher, ref 37515) 30mins at room temp. The slides were drained for a few seconds (not rinsed) and wiped around the sections with tissue paper.400ul of CDs-conjugated GFAP antibody **13a** (1:500) were then added per slide and incubated at 4°C overnight. The slides were then rinsed 3 x 5 min with TBS plus 0.05% Tween20.

#### Nuclear stain

The slides were equilibrated with 300μl buffer 2xSSC (0.3M NaCl, 0.03M sodium citrate, pH=7.0) 2x 3mins, then 150ul (500 nM) propidium iodide (Thermofisher, cat.no P3566) were added per slide, incubated at 37°C incubator for 5 mins. Afterwards, the slides were washed 6 times with buffer 2xSSC 300 μl. The slides were mounted using mounting medium fluromount-G and a coverslip was added. Clear nail polish was added to seal the edges around the coverslip.

### 4.7. Confocal Microscopy

Optical microscope images were acquired on a Leica DMIL Led Fluo microscope. Confocal microscope images were acquired on a Leica DMi8 inverted epifluorescence microscope using 405 nm and tuneable white light lasers and 63x (NA 1.4) objective at the Wolfson Imaging facility at the University of Bristol. The images were analysed using Fiji (ImageJ) software.

## Acknowledgements

The authors thank Cancer Research UK (grant number C30758/A2979). TGM thanks EPSRC BCFN EP/L016648/1/Conacyt. This research was also funded by the European Research Council (MCG), grant number ERC-COG:648239 and by the MSCA fellowship project 843720-BioNanoProbes (JRS). The authors thank Dr. Katy Jepson and the Wolfson Bioimaging Facility for her assistance in this work.

