## Supporting info for "Carbon Dot-based Fluorescent Antibody Nanoprobes as Brain Tumour Glioblastoma Diagnostics"

TABLE OF CONTENTS

1. Materials and equipment ............ S2

2. General methods ............ S3
 2.1. General protein/Abs conjugation procedure ............ S3
 2.2. Western Blot analysis . ............ S3
 2.3. Tissue staining protocol for GFAP immunofluorescence with Abs-CD 13a ............ S3

3. Chemical synthetic procedures ............ S4
 3.1. Synthesis of CD **3** ............ S4
 3.2. Synthesis of DBCO-CD **2** ............ S6
 3.3. Synthesis of azide-NHS linker **1** ............ S14

4. BSA and Abs functionalization ............ S20
 4.1 preparation of BSA derivatives **11a-f** ............ S20
 4.2. Preparation of Abs derivatives **13a-d** ............ S21

5. Microscopy images ............ S23
 5.1. Optical microscopy images ............ S23
 5.2. Confocal microscopy images ............ S23
 5.3. Clinical data for brain tumour samples ............ S28

6. Supplementary references ............ S29

### 1. Materials and equipment

**Reagents and solvents:** were purchased as reagent grade from Sigma Aldrich or ThermoFisher and used without further purification.
**Chromatography**: silica gel 60 (230-400 mesh, 0.040-0.063 mm) was purchased from E. Merck and gel filtration Sephadex G-25 was purchased from GE Healthcare. Thin Layer Chromatography (TLC) was performed on aluminium sheets coated with silica gel 60 F254 purchased from E. Merck, visualization by UV light (254 nm) and by staining with potassium permanganate or ceric molybdate solution. Extracts were concentrated in vacuo using both a Buchi rotary evaporator (bath temperatures up to 40 °C) at a pressure of either 15 mmHg (diaphragm pump) or 0.1 mmHg (oil pump), as appropriate, and a high vacuum line at room temperature.
**Dialysis purification**: Spectra Por 131096 Biotech-Grade CE Dialysis Tubing, 500-1000 MWCO, 31mm/20mm; 33ft was purchased from Cole-Parmer.
**Protein source**: BSA was purchased from ThermoFisher as Bovine Serum Albumin Standard Ampules, 2 mg/mL (23209). Polyclonal Rabbit Anti-Glial Fibrillary Acidic Protein (Concentrate) was purchased from Agilent Dako (Z033401-2). Recombinant Human GFAP protein, used in the Western Blot test was purchased from abcam (ab114149). secondary IRDye® 680RD Goat anti-Rabbit IgG used in the Western Blot test were purchased by LI-COR.
**Microwave equipment**: carbon dots were prepared using a domestic microwave oven (300 W). Tissue staining was performed on a dedicated domestic microwave oven (950 W).
**Centrifugal spin filtration**: was performed on Amicon Ultra-0.5 mL purchased from Merck using a 10K, 30K and 50K cut-off as appropriate.
**NMR equipment**: spectra were recorded on Bruker AV 400 MHz or AV 500 MHz spectrometers, using the residual solvent peaks as internal reference at 298 K. Chemical shifts are reported as parts per million and coupling constants (J) given in Hertz. Multiplicities are abbreviated as: b (broad), s (singlet), d (doublet), t (triplet), q (quartet), m (multiplet) or combinations thereof. All the assignments were confirmed by one- and two-dimensional NMR experiments (DEPT, COSY, HSQC). To confirm successful functionalisation of the CD surface with **2**, Diffusion-Ordered NMR Spectroscopy (DOSY), which probes the diffusion coefficient for each of the components of the 1H NMR spectrum, was acquired.
**Mass equipment:** high resolution mass spectra (HRMS) were obtained by the University of Bristol mass spectrometry service using electrospray ionisation (ESI) mass spectra were recorded on a Micromass LCT mass spectrometer or aVG Quattro mass spectrometer. MALDI spectra were acquired on Bruker ultrafleXtreme 2 (TOF).
**Zeta potential:** the analysis was carried out using Malvern Instruments Nano-Z ZEN 2600 and conducted in distilled $H_2O$ at a concentration of 4 mg·mL$^{-1}$.
**Plate reader equipment**: Fluorescent CDs analysis was performed BMG LABTECH CLARIO star 430-0225 plate reader.
**Gel Electrophoresis equipment:** gel electrophoresis was carried out on NuPAGE 4-12% Bis Tris Gel purchased from Invitrogen in MES buffer using a Bio-Rad 1000/500 electrophoresis power supply. Proteins were loaded at a similar concentration and stained with PageBlue™ protein staining solution.
**Western Blot equipment**: proteins were run on 12% SDS-PAGE and transferred onto membrane via Trans-Blot Turbo Transfer System (BIO-RAD) and visualized on a LI-COR Odyssey imaging system.
**Microscope equipment**: optical microscope images were acquired on a Leica DMIL Led Fluo microscope. Confocal microscope images were acquired on a Leica DMi8 inverted epifluorescence microscope using 405 nm and tuneable white light lasers and 63x (NA 1.4) objective at the Wolfson Imaging facility at the University of Bristol. The images were analysed using Fiji (ImageJ) software.

**Brain tissue samples:** samples from 13 different patients were kindly provided by the Southmead Hospital, University of Bristol, Bristol, United Kingdom.

**Brain tissue stains**: cells were stained using propidium iodide purchased from Thermofisher (P3566) for the nuclei, and compound **13a** for GFAP.

### 2. General methods

#### 2.1. General protein/Abs conjugation procedure

*Step 1 - Azide functionalization:* To a solution of protein in PBS (100 µL,36.1 µM for BSA or 100 µL, 11.1 µM Abs), different amounts of compound **1** (0.1 mg/µL in DMSO stock solution) from 0.14 to 4.81 µmol for BSA and 0.12 – 2.41 µmol for Abs, were added, respectively (See Table S1 and Table S2). The final solution was mixed in a shaker at 400 rpm for 4 h at room temperature. The product was purified via spin-filtration using 30 KDa or 50 KDa cut-off membrane for BSA or Abs respectively, at 4000 g per 20 minutes. The concentrated protein solution was diluted with 100 µL of PBS and concentrated again; this washing step was repeated two more times to remove unbound linker **1** and by-products of the reaction, furnishing a concentrated **10a-f** or **12a-d** for BSA and Abs derivatives respectively.

*Step 2 – CD-conjugation:* The concentrated **10a-f** or **12a-d** solution prepared in Step 1 was diluted with 100 µL of a PBS solution containing DBCO-CD **2** (2 mg/mL) mixed in a shaker at 400 rpm for 16 h at room temperature. The product was purified via spin-filtration using 30 KDa or 50 KDa cut-off membrane for BSA or the Abs respectively, at 4000 g per 20 minutes. The concentrated protein solution was diluted with 100 µL of PBS and concentrated again; this washing step was repeated three more times to remove the excess of **2** (4 washing steps were judged enough to remove the excess of **2** since no fluorescence was detected by the naked eye in the washing solution passing through the membrane under UV lamp in the last wash), furnishing a concentrated **11a-f** or **13a-d** solution.

#### 2.2. Western Blot analysis

Recombinant human GFAP protein was run on 12% SDS-PAGE and transferred onto membrane via Trans-Blot Turbo Transfer System (BIO-RAD). The membrane was incubated in blocking buffer (PBS, 0.1% Tween and 2% milk) for one hour and then incubated with primary anti-GFAP antibodies (Agilent, Z033401-2) or anti-GFAP-CD conjugates **13a** (results and discussion) overnight in the cold room. Membranes were washed and then incubated in secondary antibodies (IRDye® 680RD Goat anti-Rabbit IgG, LI-COR) for one hour. The membranes were washed and then visualized on a LI-COR Odyssey imaging system.

#### 2.3. Tissue staining protocol for GFAP immunofluorescence with Abs-CD 13a

The tissue sections were deparaffinized and rehydrated as follows, the sections were incubated in three washes of xylene for 2 min each, followed by two washes of 100%, 95% ethanol for 10 min each. The sections were then washed twice in distilled $H_2O$ for 5 min each.

The tissue slides were then placed in the microwaveable vessel. Tris-EDTA antigen retrieval buffer (10 mM Tris base, 1 mM EDTA solution, 0.05% Tween 20, pH 9.0) was added and placed inside the microwave, which was set to full power until the solution came to a boil. The solution was boiled for 20 min from this point and left on the bench at room temp to cool for 30mins. The slides were then washed 2 x 5 min with TBS plus 0.025% Triton X-100 with gentle agitation. The slides were blocked in Superblock buffer (Thermofisher, ref 37515) 30mins at room temp. The slides were drained for a few seconds (not rinsed) and wiped around the sections with tissue paper.400ul of CDs-conjugated GFAP antibody **13a** (1:500) were then added per slide and incubated at 4°C overnight. The slides were then rinsed 3 x 5 min with TBS plus 0.05% Tween20.

*Nuclear stain:* The slides were equilibrated with 300ul buffer 2xSSC (0.3M NaCl, 0.03M sodium citrate, pH=7.0) 2x 3mins, then 150ul (500 nM) propidium iodide/PI were added per slide, incubated at 37°C incubator for 5 mins. Afterwards, the slides were washed 6 times with buffer 2xSSC 300 µl.

The slides were mounted using mounting medium fluromount-G and a coverslip was added. Clear nail polish was added to seal the edges around the coverslip.

### 3. Synthetic procedures

#### 3.1. Synthesis of CD 3

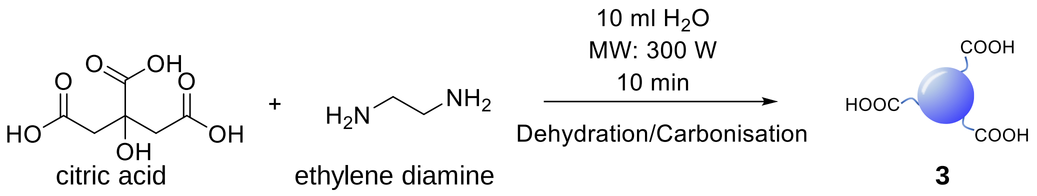


**Scheme S1.** CD synthesis.

Citric acid (1.00 g, 5.2 mmol) was dissolved in distilled $H_2O$ (10 mL) in a 250 mL conical flask. Ethylenediamine (EDA, 384 µl, 5.72 mmol) was then added to the solution and stirred for 30 min to ensure homogeneity. The conical flask was then placed in a domestic microwave 300 W (inside a fume cupboard) and the solution was reacted for 10 min. A viscous amber residue was obtained which was washed with a solution MeOH:Acetone 1:1 (4xtimes). The residue was then phase-separated by centrifugation and re-dissolved in 15 ml of distilled $H_2O$. The CD solution was dialysed in $H_2O$ using 0.5-1 KDa MWCO Biotech Cellulose Ester membrane. The concentrate CD solution was then lyophilised to yield 1.1 g of CD as an amber powder. To remove high MW components the 100 mg of CD were redissolved in $H_2O$ and filtered over Amicon Ultra spin filtration (10 KDa cut-off membrane) and liophilized furnishing 92 mg of CD as an amber powder. Procedure modified from the one reported by Mondal et al.[2]

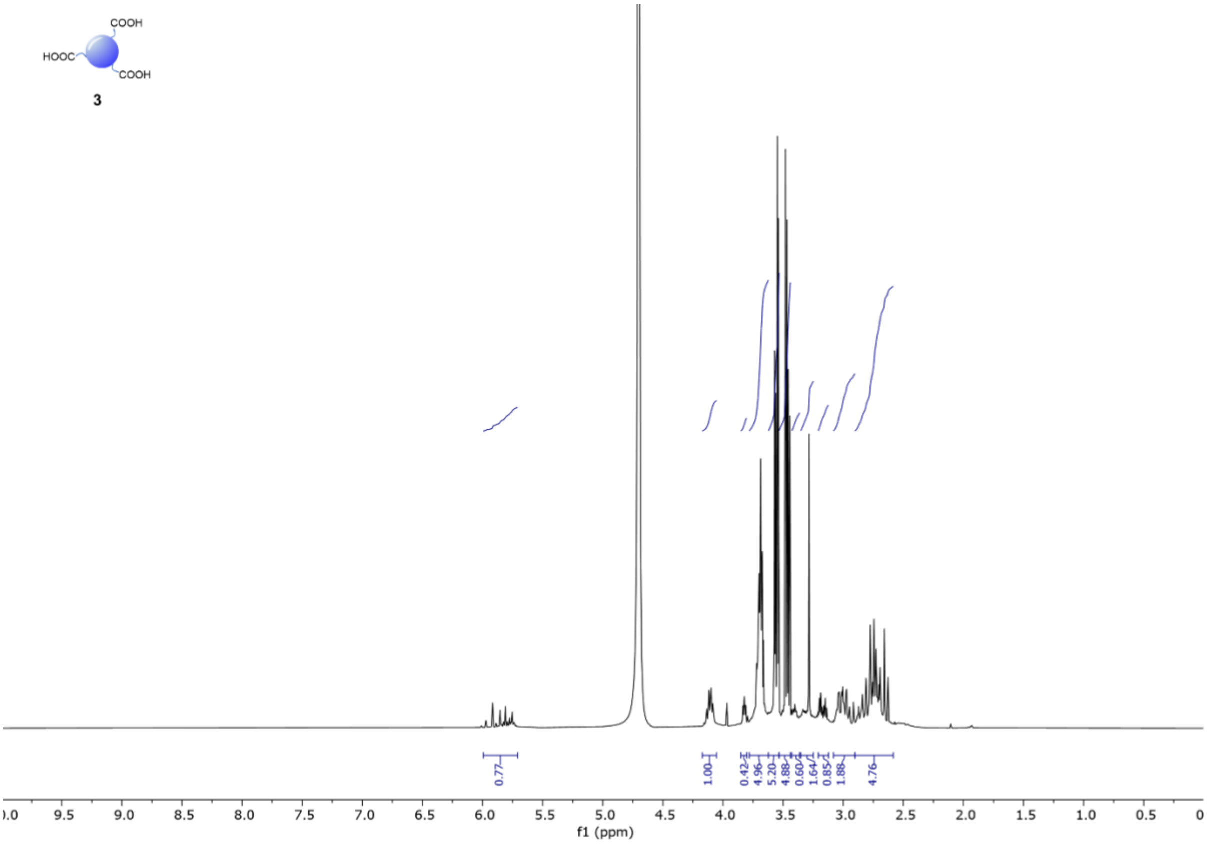


**Figure S1.** $^{1}H$ NMR spectrum (500 MHz, $D_2O$, 298 K), CDs **3**.

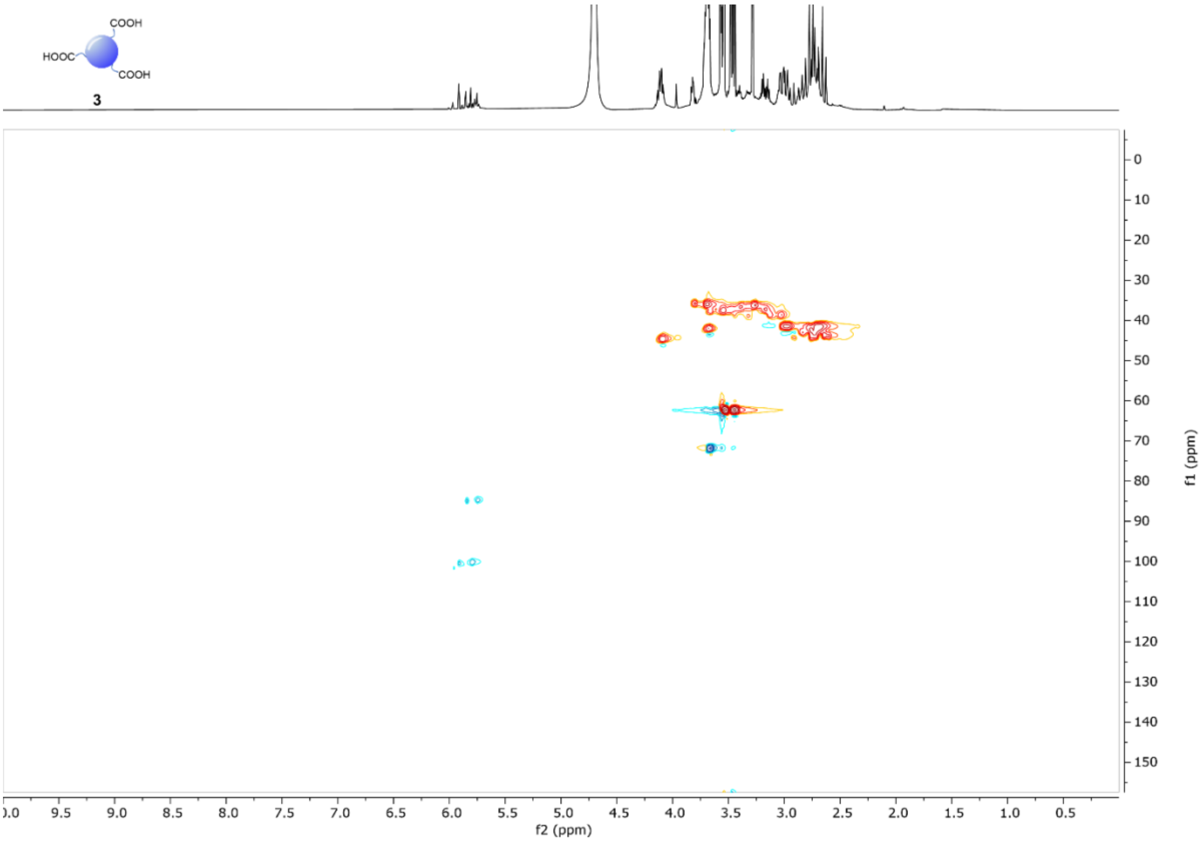


**Figure S2.** $^{1}H$-$^{13}C$ HSQC NMR spectrum ($D_2O$, 298 K), CDs **3**.

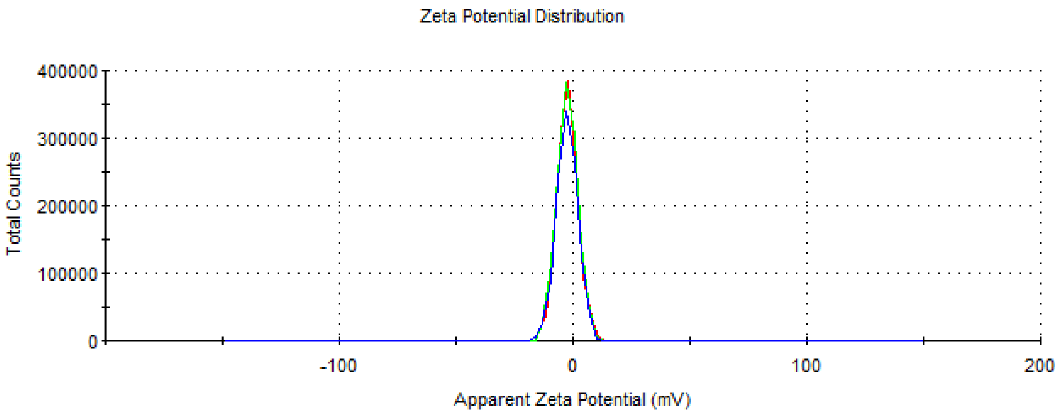


-3.09 ± 0.12 mV

**Figure S3**. Zeta-potential distribution of CD 3. Each line represents individual measurements of the same sample, which were used to obtain an average value.

#### 3.2. Synthesis of DBCO-CD 2

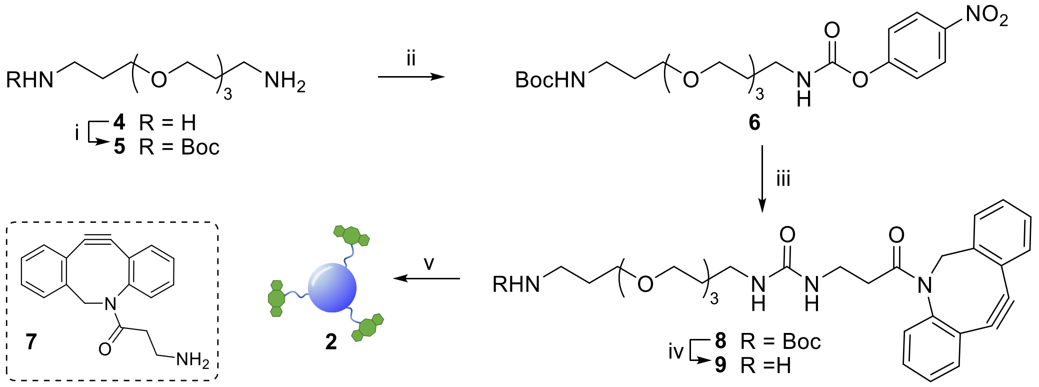


**Scheme S2**. *Reagents and conditions*: i) $Boc_2O$, DCM, 4 h, 0 °C to rt, 99 %; ii) 4-nitrophenyl chloroformate, Py, DCM, 3 h, 0 °C to rt, 87 %; iii) **7**, Py, DIPEA, DMF, 63 %; iv) TFA, DCM, 1.5 h, rt, 86 %; v) **3**, HATU, DIPEA, DMF, 5 h, rt.

**Compound 5**

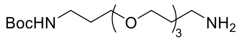


To a stirred solution of 4,7,10-Trioxa-1,13-tridecanediamine **4** (4.40 g, 20.07 mmol) in DCM (20 mL), a solution of $Boc_2O$ (435 mg. 2.00 mmol) in DCM (10 mL) was added dropwise over 1 h at 0 °C. Once the addition was completed, the mixture was allowed to stir for further 3 h at room temperature. The solution was then diluted with DCM (200 mL) and washed with brine (3 x 50 mL), and $H_2O$ (1 x 50 mL). The organic phase was dried with anhydrous $MgSO_4$, filtered and concentrated under reduced pressure furnish **5** (635 mg, 99 % yield) as a transparent liquid. **$^1$H NMR** (400 MHz, Chloroform-*d*) δ 3.66 – 3.48 (m, 12H, $OCH_2$), 3.20 (q, *J* = 6.3 Hz, 2H, $CH_2NHBoc$), 2.79 (t, *J* = 6.7 Hz, 2H, $CH_2NH_2$), 1.73 (h, *J* = 6.4 Hz, 4H, $CH_2CH_2CH_2$), 1.42 (s, 9H, $CH_3^{Boc}$). NMR data are in agreement to those reported in the literature.[1]

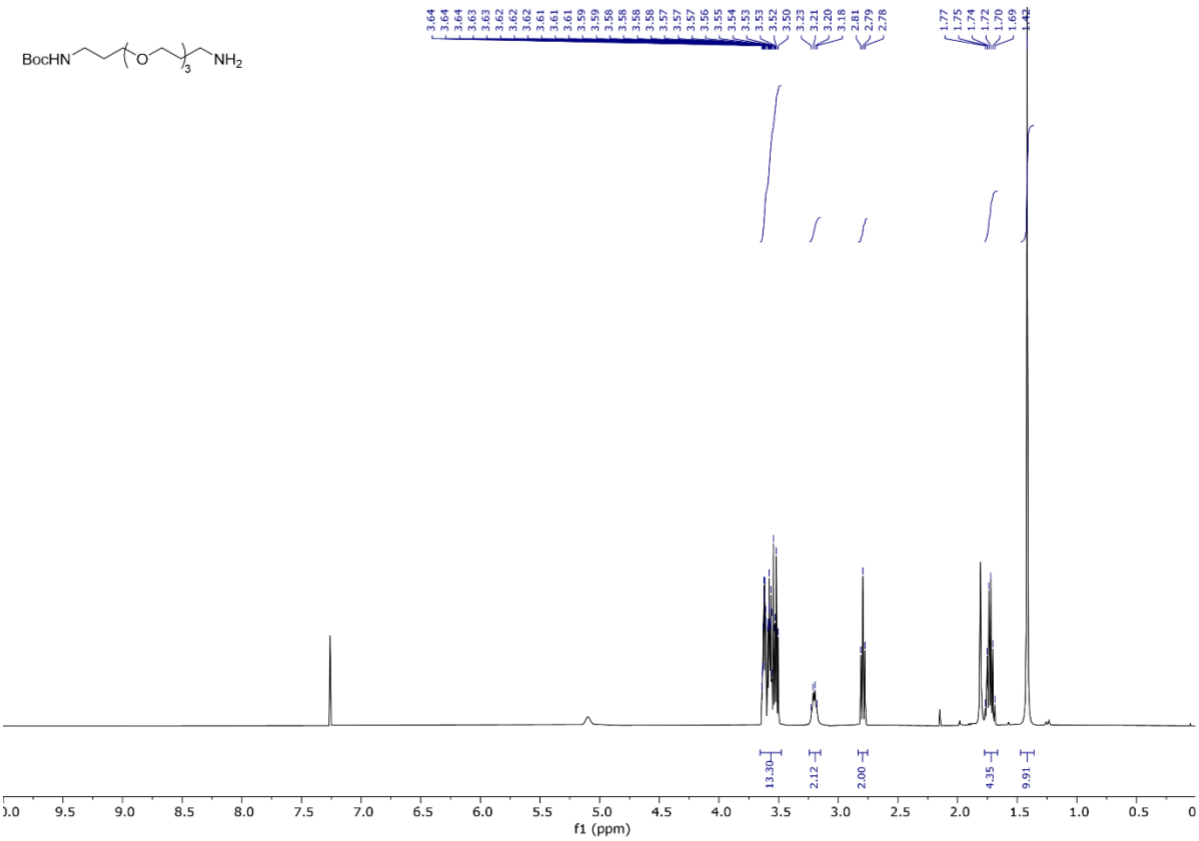


**Figure S4.** $^1$H NMR spectrum (400 MHz, $CDCl_3$), compound **5**.

**Compound 6**

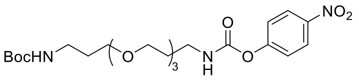


To a stirred solution of 4-nitrophenyl chloroformate (2.98 g, 14.79 mmol) and Py (1 mL, 12.33 mmol) in anhydrous DCM (70 mL) at 0 °C, a solution of **5** (1.58 g, 4.93 mmol) in dry DCM (20 mL), was added over 1 h at 0 °C. Once the addition was completed, the solution was stirred for further 2 h at room temperature. The reaction was quenched by the addition of saturated aq. $NH_4Cl$ solution (50 mL) and the mixture was extracted with DCM (3 x 50 mL). The combined organic phases were dried with anhydrous $MgSO_4$, filtered and concentrated under reduced pressure. The residue was purified by column chromatography on silica gel (Hex/EtOAc 1:0 to 3:7, v/v) furnishing **6** (2.08 g, 87 % yield) as a transparent syrup. **$^1$H NMR** (500 MHz, Chloroform-*d*) δ 8.27 – 8.20 (m, 2H, Ar), 7.35 – 7.27 (m, 2H, Ar), 6.12 (s, 1H, NH), 4.89 (s, 1H, NH), 3.69 – 3.62 (m, 8H, $OCH_2$), 3.59 (dd, $J$ = 5.8, 3.5 Hz, 2H, $OCH_2$), 3.51 (t, $J$ = 6.0 Hz, 2H, $OCH_2$), 3.42 (q, $J$ = 6.0 Hz, 2H, $NCH_2$), 3.21 (q, $J$ = 6.5 Hz, 2H, $NCH_2$), 1.87 (h, $J$ = 6.0 Hz, 2H, $CH_2CH_2CH_2$), 1.77 – 1.69 (m, 2H, $CH_2CH_2CH_2$), 1.43 (s, 9H, $CH_3$). **$^{13}$C NMR** (126 MHz, $CDCl_3$) δ 156.3, 156.2, 153.4, 144.8, 125.2, 122.1, 79.2, 70.7, 70.7, 70.4, 70.3, 70.1, 69.7, 40.2, 38.6, 29.8, 29.1, 28.6. **HRMS (ESI)** m/z: Calcd for $C_{22}H_{35}N_3O_9Na$ $(M+Na)^+$ 508.2265, found 508.2279.

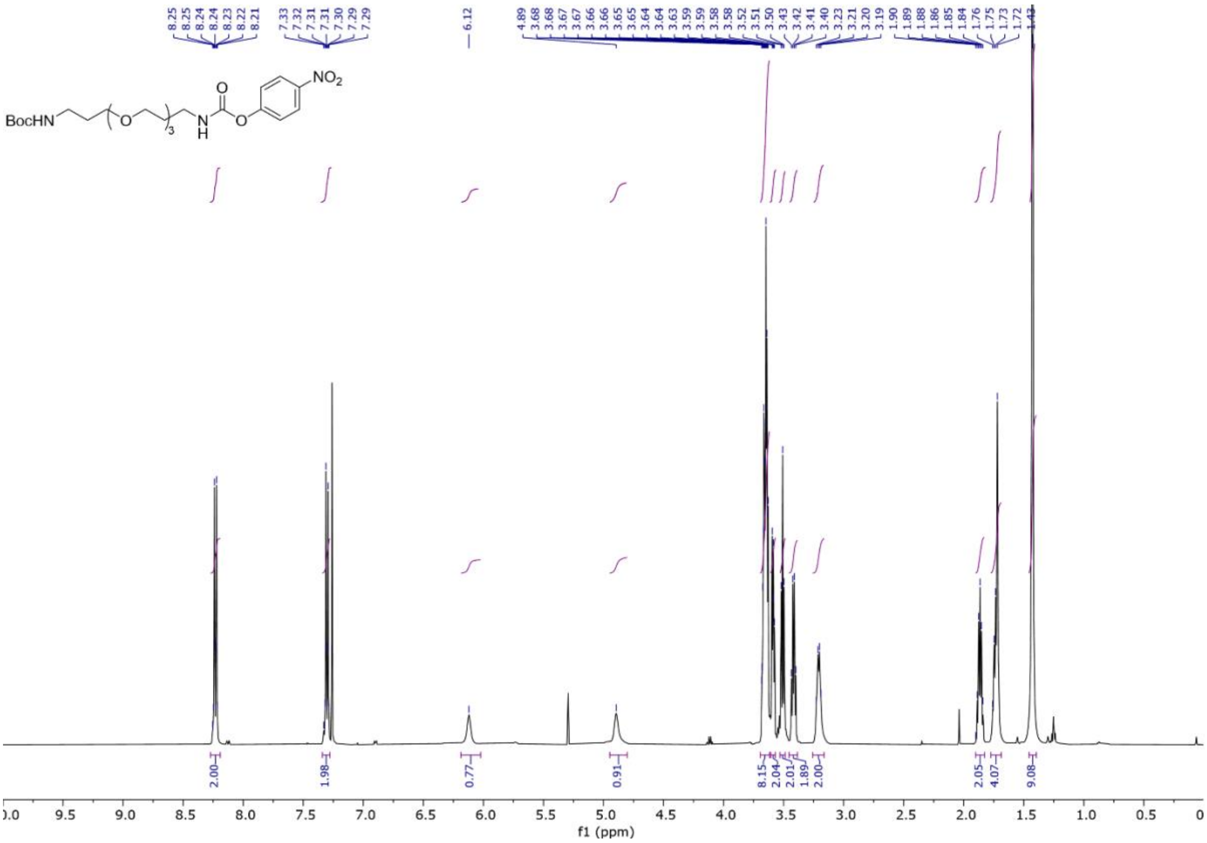


**Figure S5.** $^{1}$H NMR spectrum (500 MHz, $CDCl_3$), compound **6**.

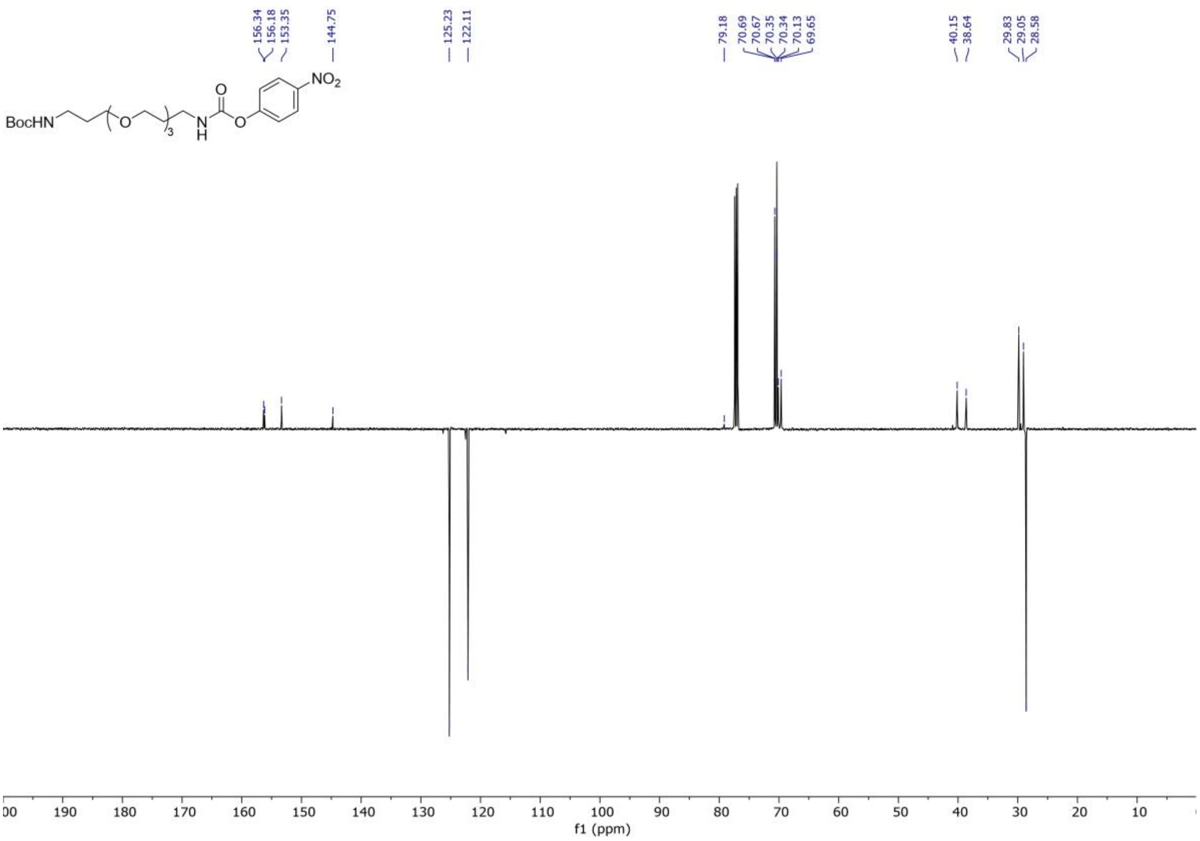


**Figure S6.** $^{13}$C APT NMR spectrum (126 MHz, $CDCl_3$), compound **6**.

**Compound 8**

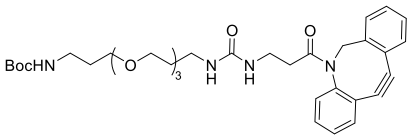


To a stirred solution of **6** (81 mg, 0.29 mmol) and Py (0.5 mL, 6.21 mmol) in anhydrous DCM (5 mL), a solution of DBCO-amine **7** (286 mg, 0.59 mmol) in anhydrous DCM (2 mL) was added dropwise at room temperature, followed by the addition of DIPEA (153 µL, 0.88 mmol) at room temperature. The solution was stirred for 4 h at room temperature, then diluted with DCM (100 mL), washed wit saturated aq. $NH_4Cl$ solution (2 x 50 mL) and brine (1 x 50 mL). The organic phase was dried with anhydrous $MgSO_4$, filtered, and concentrated under reduced pressure. The residue was purified by column chromatography on silica gel (EtOAc/MeOH 1:0 to 9:1, v/v) furnishing **8** (116 mg, 63 % yield) as a transparent oil. **$^1$H NMR** (500 MHz, $CDCl_3$) δ 7.66 (d, *J* = 7.6 Hz, 1H, Ar), 7.42 – 7.23 (m, 7H, Ar), 5.12 (d, *J* = 13.9 Hz, 1H, $CH_{2a}^{DBCO}$), 5.05 (s, 1H, NH), 5.00 (d, *J* = 6.3 Hz, 1H, NH), 4.86 (s, 1H, NH), 3.69 – 3.47 (m, 13H, $CH_{2b}^{DBCO}$, $OCH_2$), 3.30 – 3.10 (m, 6H, $NCH_2$), 2.56 – 2.48 (m, 1H, $COCH_2$), 1.98 – 1.84 (m, 1H, $COCH_2$), 1.79 – 1.63 (m, 4H, $NCH_2CH_2CH_2$), 1.42 (s, 9H, $CH_3$). **$^{13}$C NMR** (126 MHz, $CDCl_3$) δ 172.5, 158.3, 156.1, 151.2, 148.1, 132.1, 129.2, 128.6, 128.2, 128.2, 127.7, 127.1, 125.5, 123.1, 122.5, 114.7, 107.9, 70.5, 70.4, 70.1, 69.8, 69.5, 69.4, 55.5, 38.4, 36.1, 35.6, 29.6, 29.5, 28.4. **HRMS (ESI)** m/z: Calcd for $C_{34}H_{47}N_4O_7Na$ $(M+Na)^+$ 645.3259, found 645.3236.

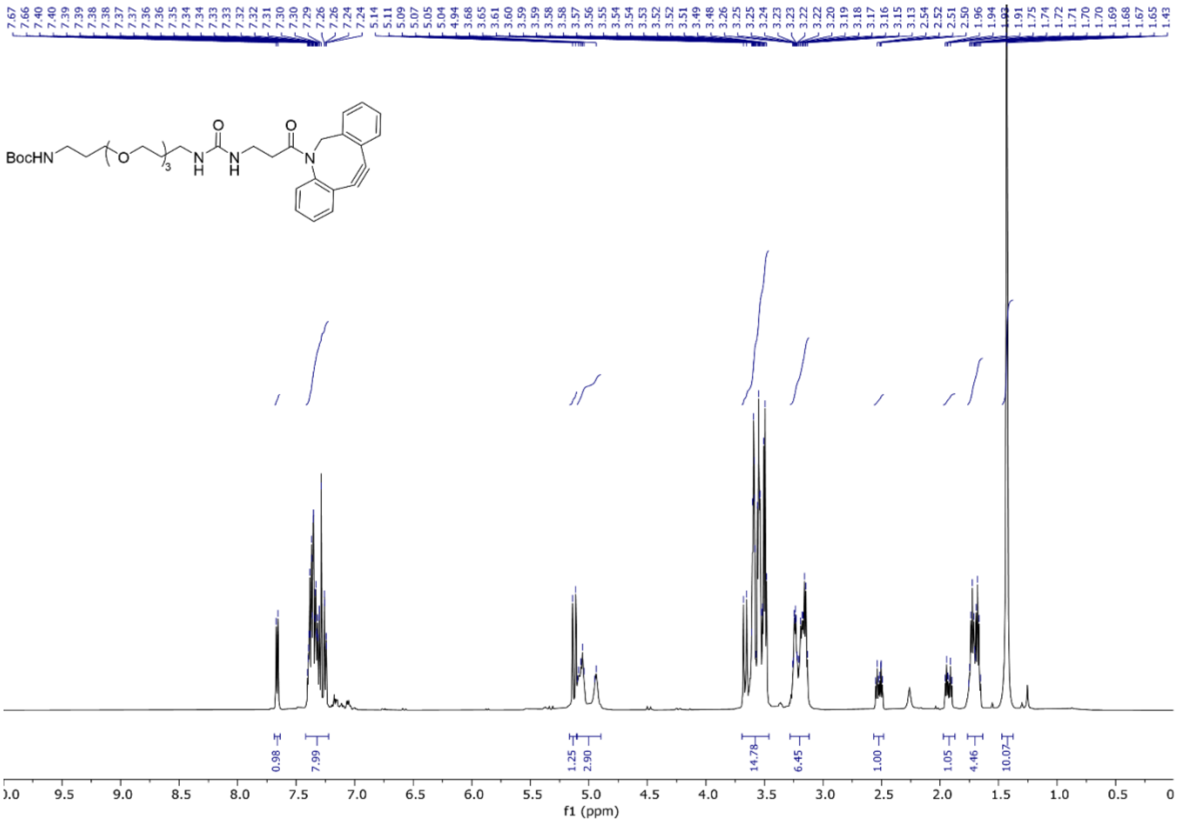


**Figure S7**. $^1$H NMR spectrum (500 MHz, $CDCl_3$), compound **8**.

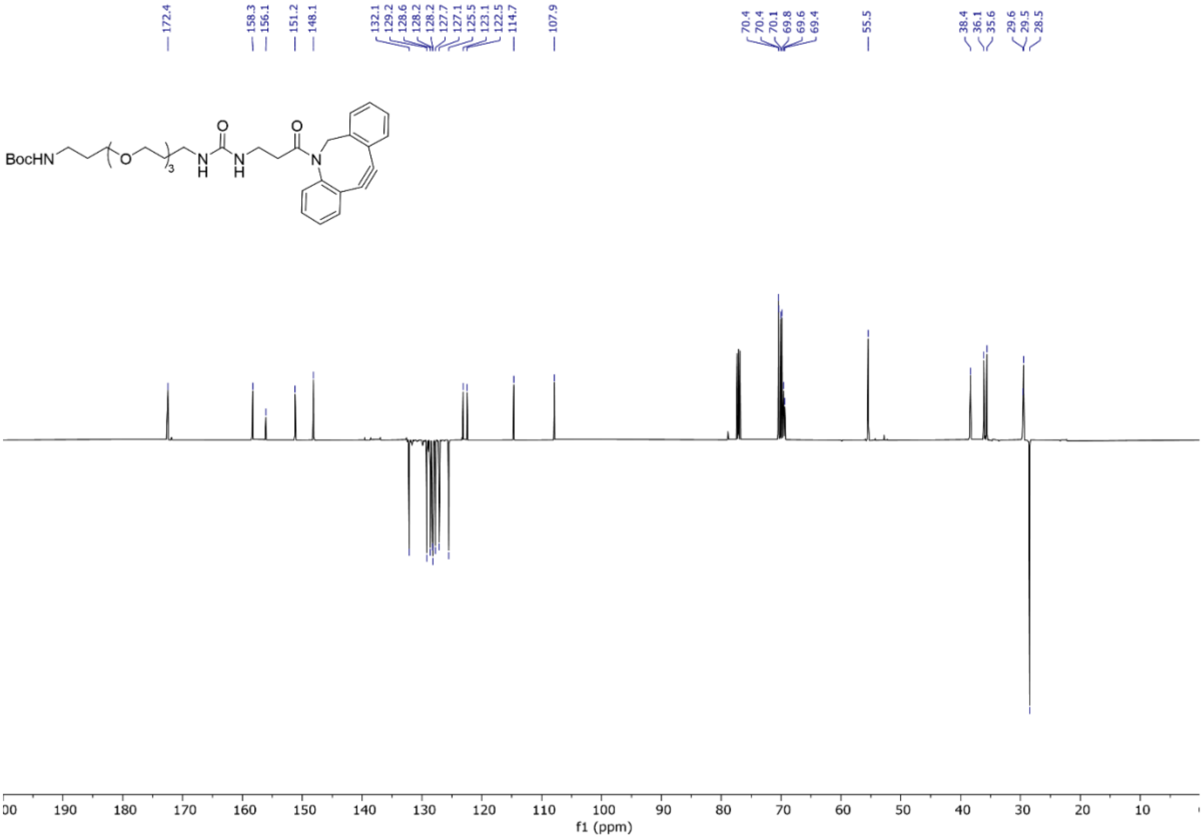


**Figure S8.** $^{13}C$ APT NMR spectrum (126 MHz, $CDCl_3$), compound **8**.

**Compound 9**

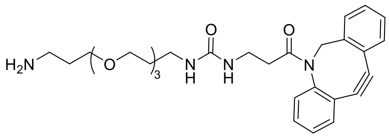


Compound **8** (116 mg, 0.19 mmol) was dissolved in a DCM/TFA solution (7 mL, 95:5, v/v) and stirred for 1.5 h at room temperature. The reaction was concentrated under reduced pressure and the residue was purified by column chromatography on silica gel ($CHCl_3$/MeOH 1:0 to 95:5, containing a 0.5% of 35% aq. $HN_4OH$ solution v/v/v) furnishing **9** (84 mg, 86 % yield) as a pale brown oil. **$^1$H NMR** (500 MHz, $D_2O^{25°C}$) δ 7.63 (d, *J* = 7.7 Hz, 1H, Ar), 7.46 – 7.35 (m, 6H, Ar), 7.29 – 7.23 (m, 1H, Ar), 5.02 (d, *J* = 14.5 Hz, 1H, $CH_{2a}^{DBCO}$), 3.72 – 3.61 (m, 11H, $CH_{2b}^{DBCO}$, $OCH_2$), 3.52 (t, *J* = 6.4 Hz, 2H, $OCH_2$), 3.14 – 2.95 (m, 6H, $NCH_2$), 2.32 – 2.16 (m, 2H, $CH_2CO$), 1.94 (dt, *J* = 13.5, 6.4 Hz, 2H, $NCH_2CH_2CH_2$), 1.67 (p, *J* = 6.6 Hz, 2H, $NCH_2CH_2CH_2$). **$^{13}$C NMR** (126 MHz, $D_2O^{25°C}$) δ 174.3, 159.7, 150.6, 147.7, 131.9, 129.1, 129.1, 128.9, 128.5, 128.1, 127.0, 125.7, 122.4, 121.6, 114.3, 107.8, 69.6, 69.5, 69.4, 69.3, 68.5, 68.3, 55.5, 37.6, 36.8, 36.3, 34.7, 29.0, 26.5. **HRMS (ESI)** m/z: Calcd for $C_{29}H_{39}N_4O_5$ $(M+H)^+$ 523.2915, found 523.2927.

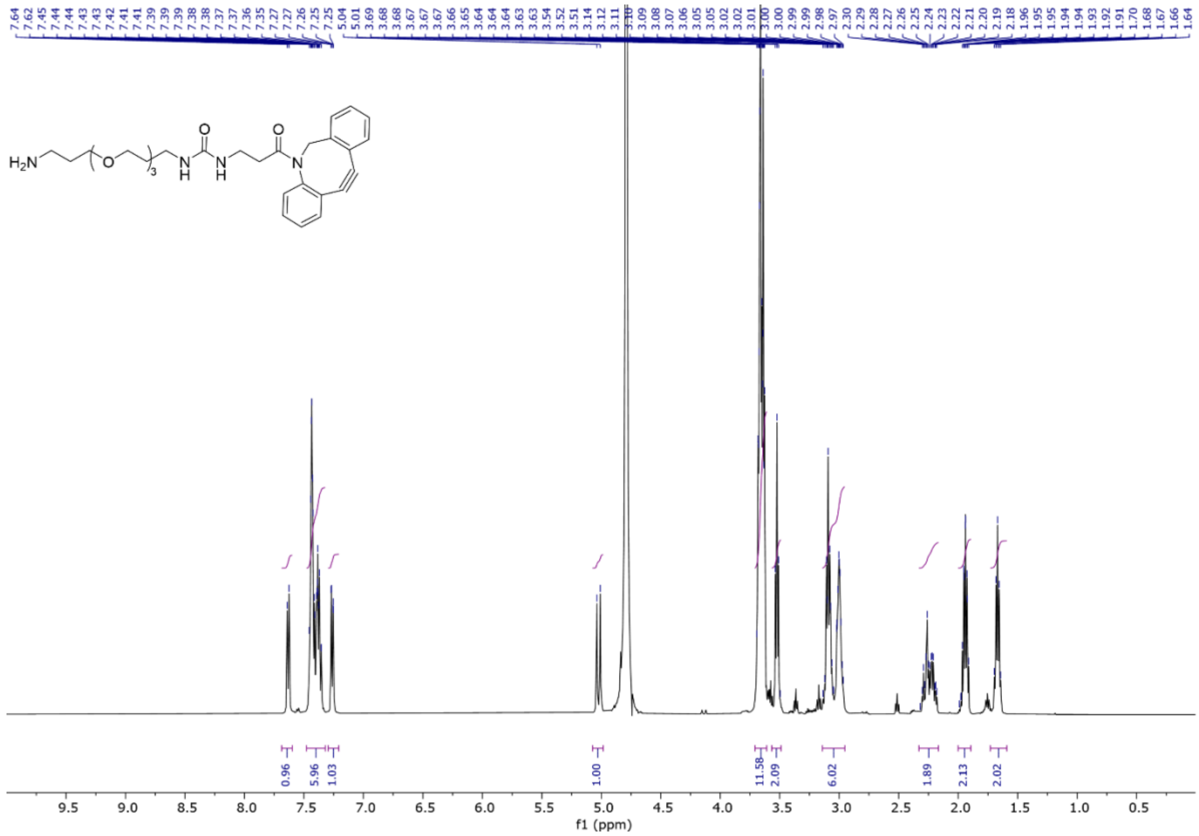

**Figure S9.** $^{1}H$ NMR spectrum (500 MHz, $CDCl_3$), compound **9**.

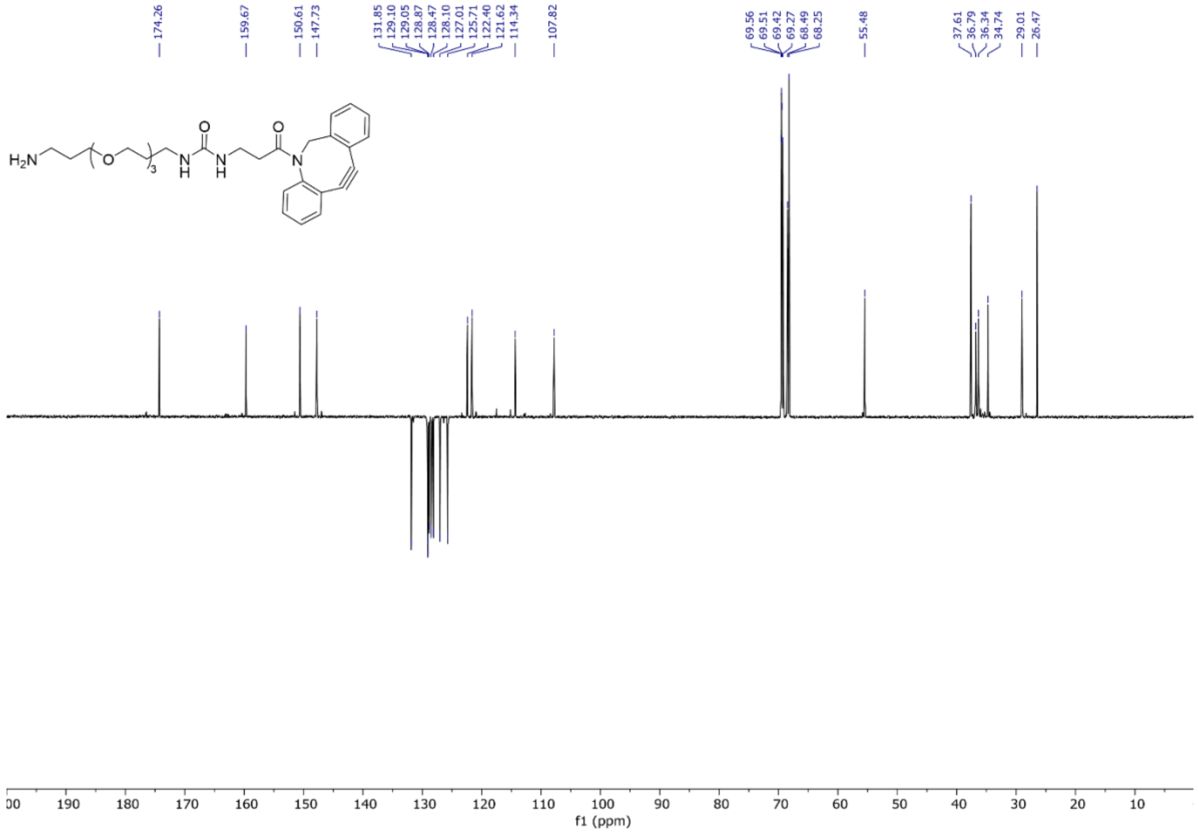

**Figure S10.** $^{13}C$ APT NMR spectrum (126 MHz, $CDCl_3$), compound **9**.

**DBCO-CD 2**

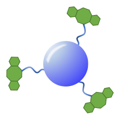

To a stirred solution of CDs **3** (18.4 mg) in dry DMF (1.84 mL), HATU (13.4 mg, 0.035 mmol) and DIPEA (6.1 µL, 0.035 mmol) were added and the solution was allowed to stir for further 15 minutes at room temperature. A solution of **9** (9.2 mg, 0.018 mmol) in dry DMF (0.5 mL) was added and the solution was stirred at room temperature for 5 h. $H_2O$ (0.5 mL) was then added to quench the reaction and the solution was stirred for further 10 minutes at room temperature and concentrated under reduced pressure. The residue was redissolved in aq. 0.1 M NaOH solution (3 mL) and stirred for 1 h at room temperature. The pH was neutralized with the addition of aq. HCl 1M solution (0.15 mL), diluted with $H_2O$ (20 mL), washed with $Et_2O$ (5 x 10 mL), and the water phase was concentrated under reduced pressure. The residue was purified via 1 KDa cut-off dialysis membrane against water, changing the water bath 3 times over a 24 h period. The purified solution was then freeze-dried furnishing **2** (10.2 mg) as a pale yellow solid.**$^1$H NMR** (500 MHz, $D_2O$, 25 °C) characteristic resonances δ ppm = 7.77 – 7.06 (m, Ar), 5.14 – 5.04 (m, $CH_2^{DBCO}$a) 4.29 – 2.50 (PEG linker and $CH_2^{DBCO}$b). **$^1$H-$^{13}$C NMR HSQC** (126 MHz, $D_2O$ 25 °C) characteristic resonances δ ppm = 131.6 (Ar), 127.1 (Ar), 129.1 (Ar), 125.8 (Ar), 55.5 ($CH_2^{DBCO}$), 44.5, 41.9, 69.3, 68.4, 39.0, 38.4, 36.1, 36.4, 36.4, 43.8, 44.2, 36.4, 36.4, 34.5, 28.3, 19.6, 0.6.

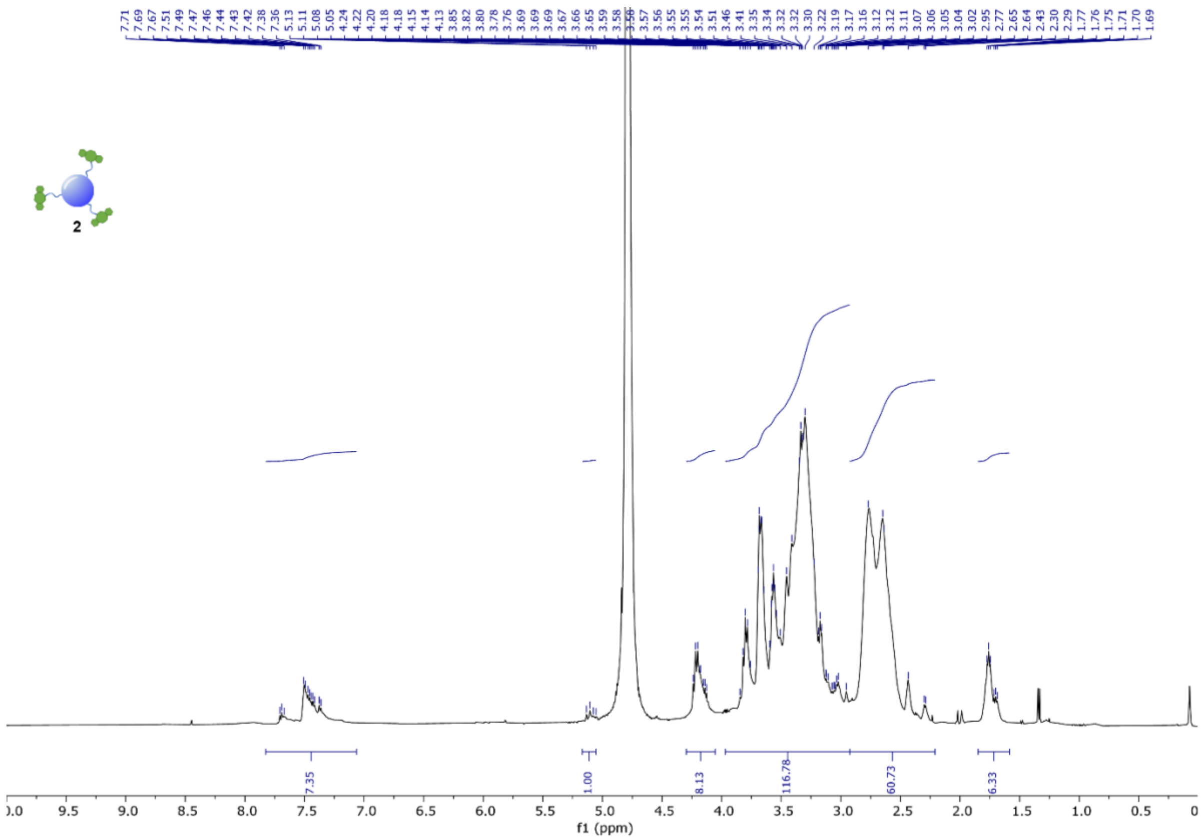


**Figure S11.** $^1$H NMR spectrum (500 MHz, $D_2O$, 298 K), DBCO-CD **2**.

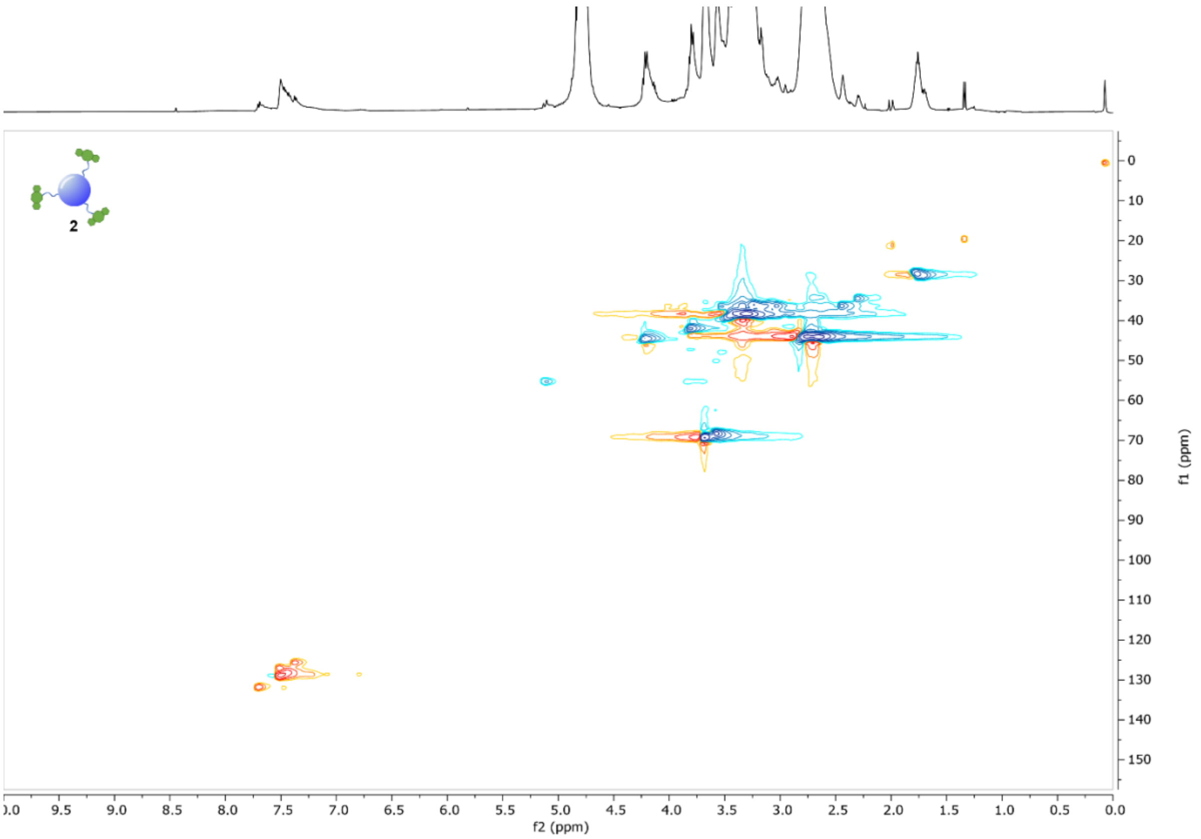


**Figure S12.** $^{1}H$-$^{13}C$ HSQC NMR spectrum ($D_2O$, 298 K), DBCO-CD **2**.

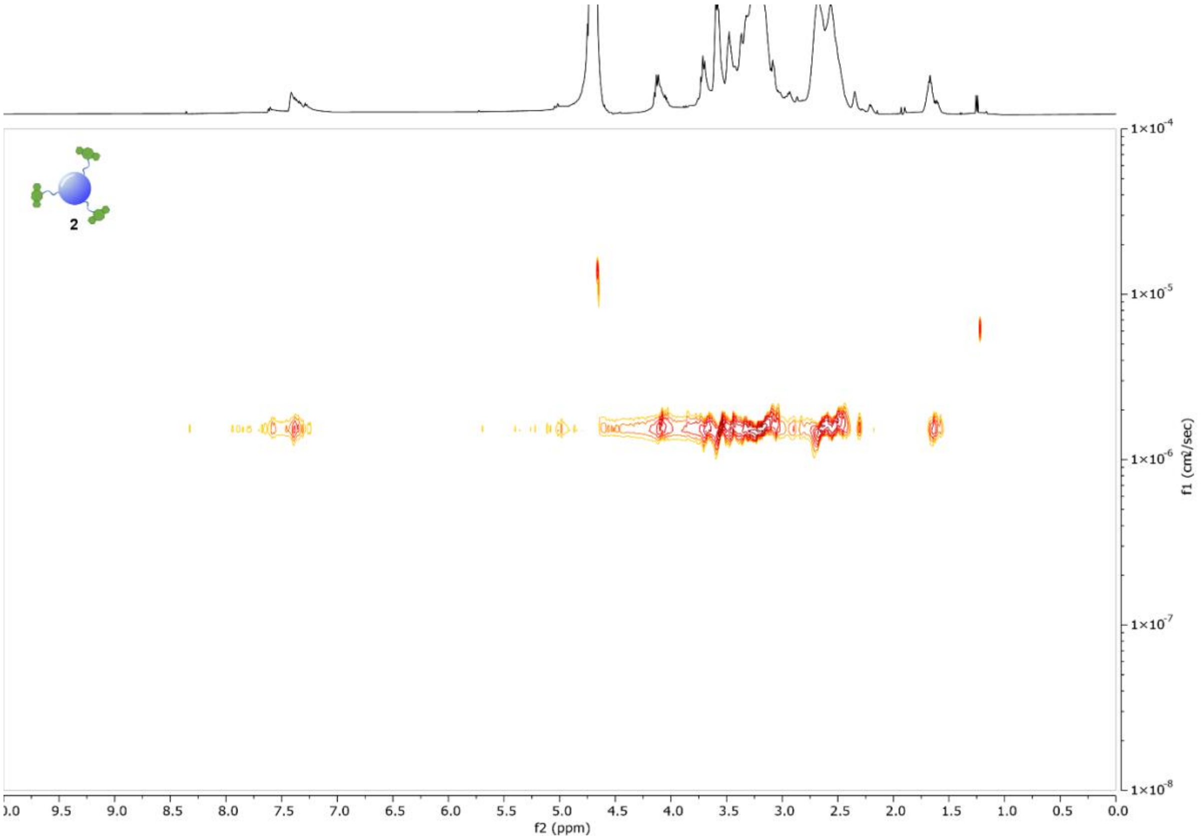


**Figure S13.** $^{1}H$ DOSY NMR spectrum (500 MHz, $D_2O$, 298 K), DBCO-CD **2**.

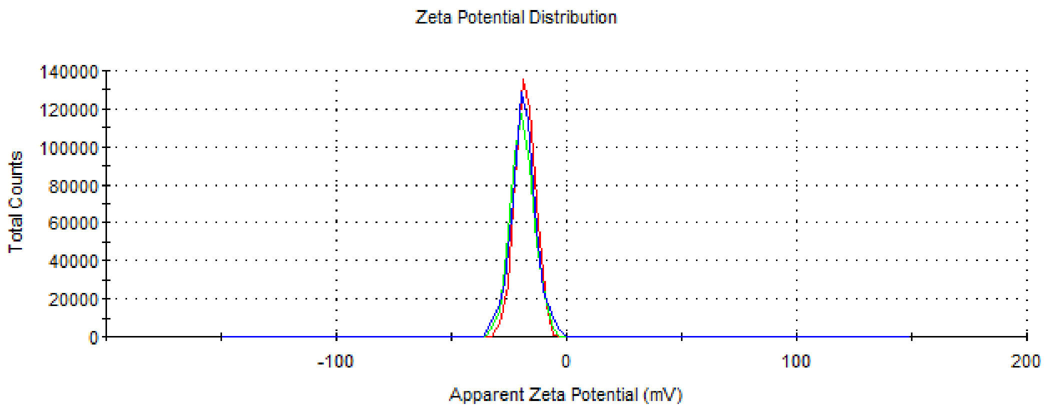


-18.0 ± 4.3 mV

**Figure S14.** Zeta-potential distribution of CD-DBCO 2. Each line represents individual measurements of the same sample, which were used to obtain an average value.

#### 3.3. Synthesis of azide-NHS linker 1

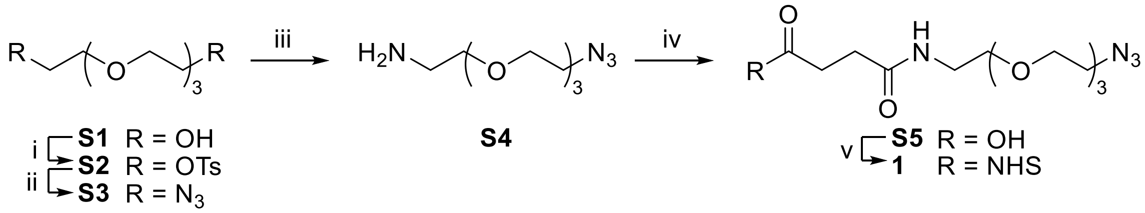


**Scheme S3.** *Reagents and conditions*: i) TsCl, $Et_3N$, DCM, 16h, rt, quant.; ii) $NaN_3$, DMF, 16 h, 40 °C, 76 %; iii) $PPh_3$, 5 % aq. HCl, $Et_2O$, THF, 16h, rt, 74%; iv) Succinic anhydride, DIPEA, DCM, 3 h, rt, 52 %; v) NHS, EDC·HCl, DCM, 2h, rt, 93 %.

#### Compound S2

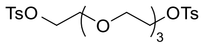

To a stirred solution of tetraethylene glycol **S1** (11.25 g, 57.92 mmol) in DCM (80 mL) triethylamine (24.2 mL, 176.0 mmol) and tosyl chloride (33.2 g, 174.0 mmol) were added with immediate precipitation triethylammonium chloride as a white sold. The suspension was stirred 16 h, at room temperature. Filtered over a sintered funnel to remove solids and the solid was washed with DCM (2 x 20 mL). The liquid filtrate was diluted with DCM (400 mL) washed with $H_2O$ (3 x 100 mL), and brine (1 x 100 mL). The organic phase was dried with anhydrous $MgSO_4$, filtered and concentrated under reduced pressure. The residue was purified by column chromatography on silica gel (Hex/EtOAc 1:0 to 0:1, v/v) to furnish **S2** (29.1 g, quant.) as an orange syrup. $^1H$ NMR (400 MHz, $CDCl_3$) δ 7.83 – 7.75 (m, 4H, Ar), 7.37 – 7.29 (m, 4H, Ar), 4.18 – 4.10 (m, 4H, $TsOCH_2$), 3.71 – 3.64 (m, 4H $TsOCH_2CH_2$), 2.44 (s, 6H, $CH_3$). NMR data are in agreement to those reported in the literature.[1]

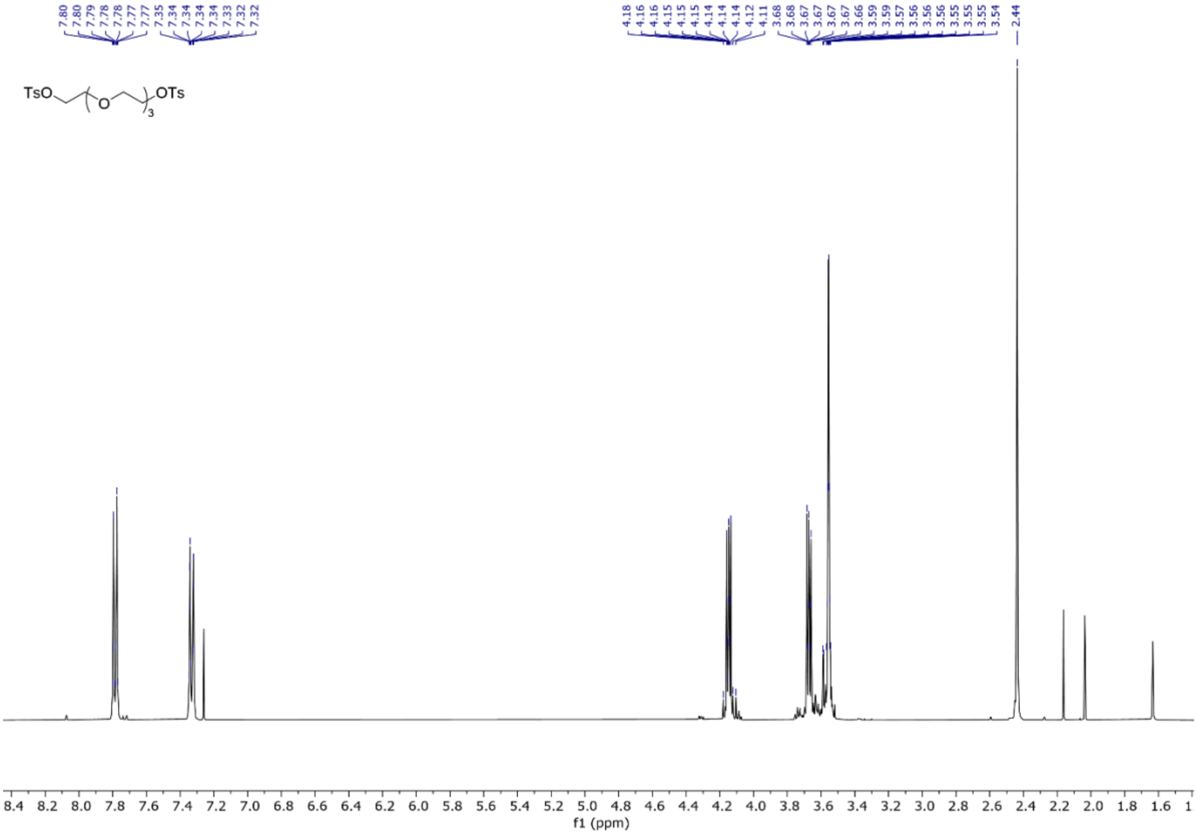


**Figure S15**. $^1$H NMR spectrum (400 MHz, $CDCl_3$), compound **S2**.

**Compound S3.**

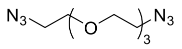


To a stirred solution of **S2** (25.9 g, 51.5 mmol) in DMF (115 mL), $NaN_3$ (15.0 g, 230.6 mmol) was added and the mixture was stirred for 16 h at 40 °C. The mixture was concentrated under reduced pressure and the residue was resuspended in EtOAc (300 mL) and washed with $H_2O$ (3 x 100 mL), and brine (1 x 100 mL). The organic phase was dried with anhydrous $MgSO_4$, filtered and concentrated under reduced pressure. The residue was purified by column chromatography on silica gel (Hex/EtOAc 1:0 to 7:3, v/v) to furnish **S3** (10.7 g, 76 % yield) as a pale yellow liquid. $^1$H NMR (400 MHz, Chloroform-*d*) δ 3.69 – 3.62 (m, 12H, $OCH_2$), 3.37 (t, *J* = 5.1 Hz, 4H, $N_3CH_2$). NMR data are in agreement to those reported in the literature.[3]

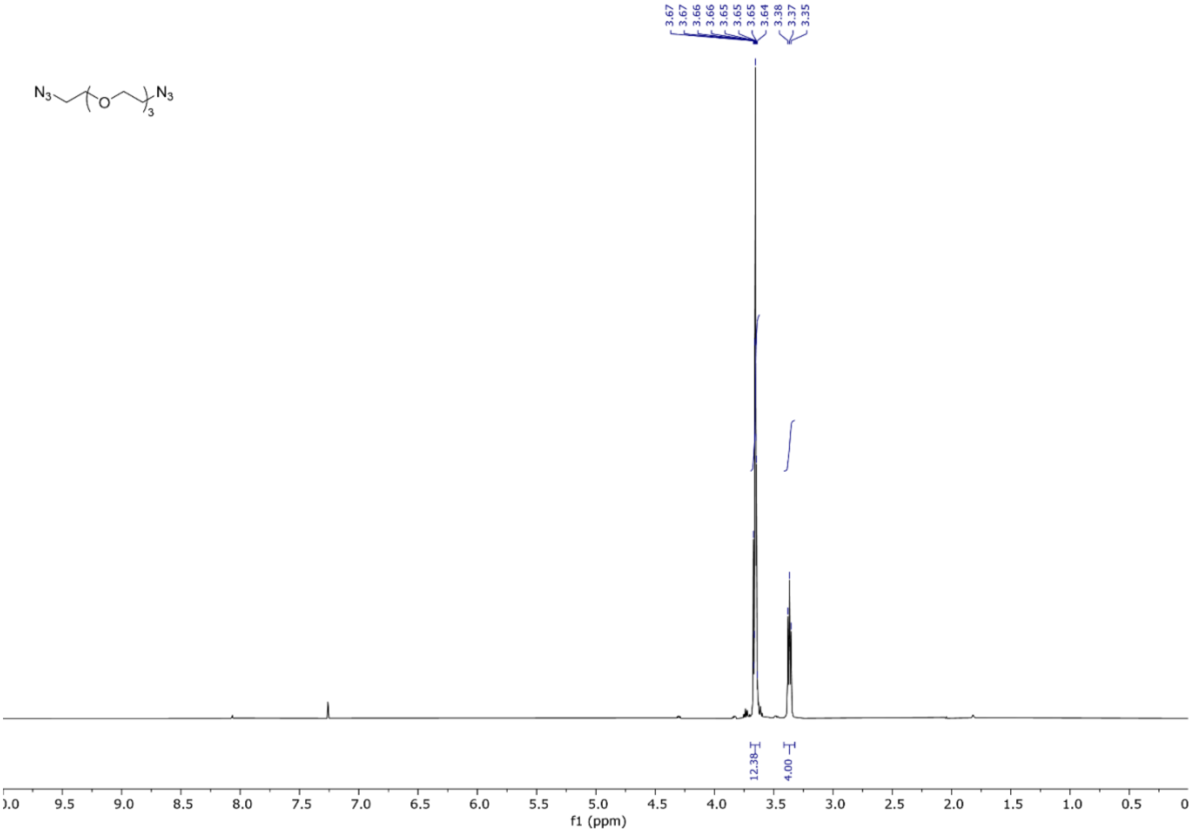


**Figure S16.** $^1$H NMR spectrum (400 MHz, $CDCl_3$), compound **S3**.

Compound **S4**.

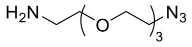


To a stirred solution of **S3** (0.96 g, 3.93 mmol) in THF (5 mL) and aqueous 5 % HCl (12 mL) a solution of $PPh_3$(0.927 g, 3.54 mmol) in $Et_2O$ (18 mL) was added over 30 minutes with a dropping funnel and vigorously stirred for 16 h at room temperature. The phases were separated and the water layer was washed with DCM (3 x 10 mL) and the organic layer was discarted. The pH of the aqueous phase was adjusted to c.ca 10 using KOH and extracted with DCM (4 x 20 mL). The combined organic layers were dried with anhydrous $MgSO_4$, filtered and concentrated under reduced pressure furnishing **S4** (0.632 g, 74 % yield) as a transparent oil. **$^1$H NMR** (400 MHz, $CDCl_3$) δ 3.70 – 3.60 (m, 10H, $OCH_2$), 3.50 (td, *J* = 5.3, 1.3 Hz, 2H, $OCH_2$), 3.42 – 3.35 (m, 2H, $CH_2NH_2$), 2.86 (td, *J* = 5.3, 1.4 Hz, 2H, $CH_2N_3$), 1.49 (s, 2H, $NH_2$). NMR data are in agreement to those reported in the literature.[3]

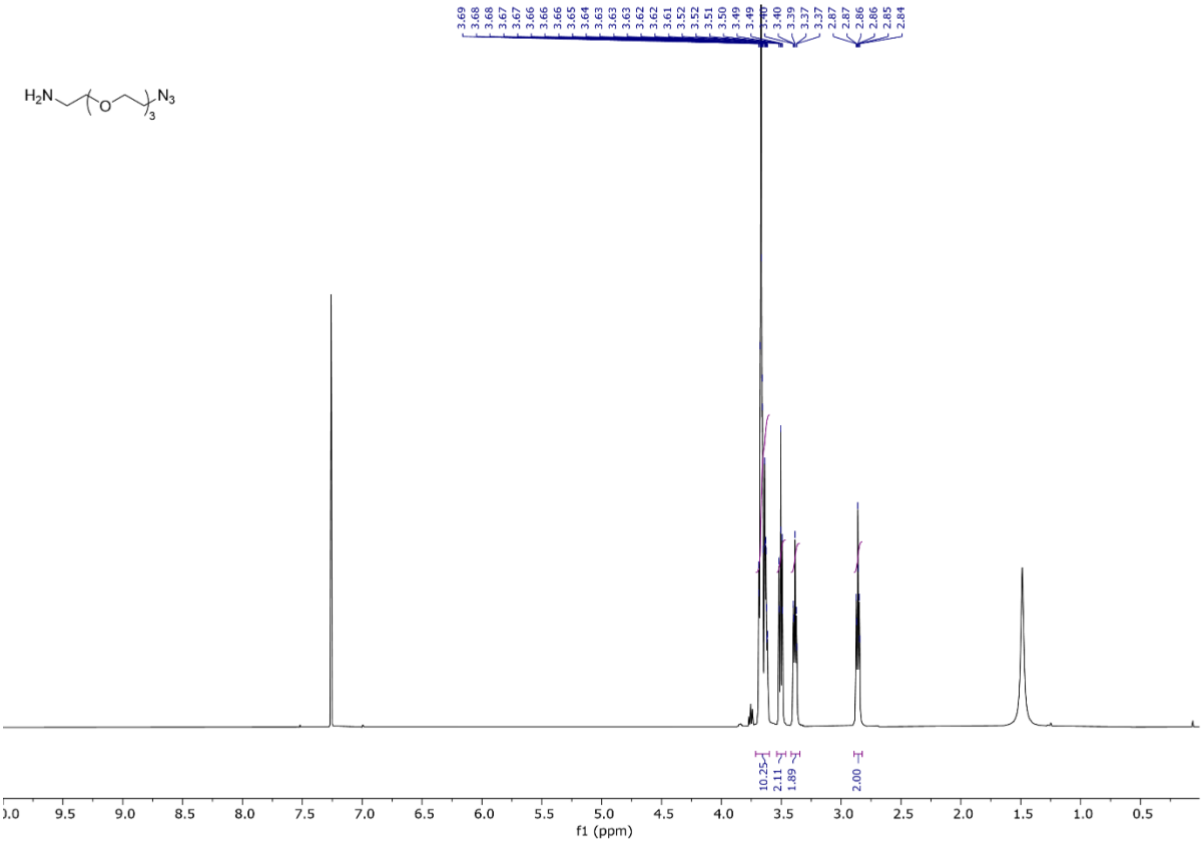


**Figure S17.** $^1$H NMR spectrum (400 MHz, $CDCl_3$), compound **S4**.

**Compound S5**.

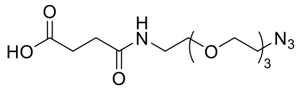


To a stirred solution of **S4** (632 mg, 2.90 mmol) in DCM (15 mL), succinic anhydride (580 mg, 5.79 mmol) and DIPEA (1.26 mL, 7.24 mmol) were added at room temperature. The solution was stirred for 3 h at room temperature and then quenched by the addition of saturated aq. $NaHCO_3$solution. The pH of the final solution was then acidified to pH < 1 with aq. 5M HCl, diluted with water (20 mL) and extracted with DCM (4 x 20 mL). The combined organic layers were dried with anhydrous $MgSO_4$, filtered and concentrated under reduced pressure. The residue was purified by column chromatography on silica gel ($CHCl_3$/MeOH 1:0 to 97:3, containing 0.3 % pf AcOH, v/v/v) furnishing **S5** (0.48 g, 52 % yield) as a pale brown oil. $^1$H NMR (500 MHz, $CDCl_3$) δ 3.73 – 3.60 (m, 10H, $OCH_2$), 3.55 (t, *J* = 5.0 Hz, 2H, $OCH_2CH_2NH$), 3.45 (t, *J* = 5.2 Hz, 2H, $OCH_2CH_2NH$), 3.40 (t, *J* = 5.0 Hz, 2H, $N_3CH_2$), 2.67 (dd, *J* = 7.6, 5.5 Hz, 2H, $CH_2CH_2$), 2.53 (dd, *J* = 7.5, 5.6 Hz, 2H, $CH_2CH_2$). NMR data are in agreement to those reported in the literature.[1]

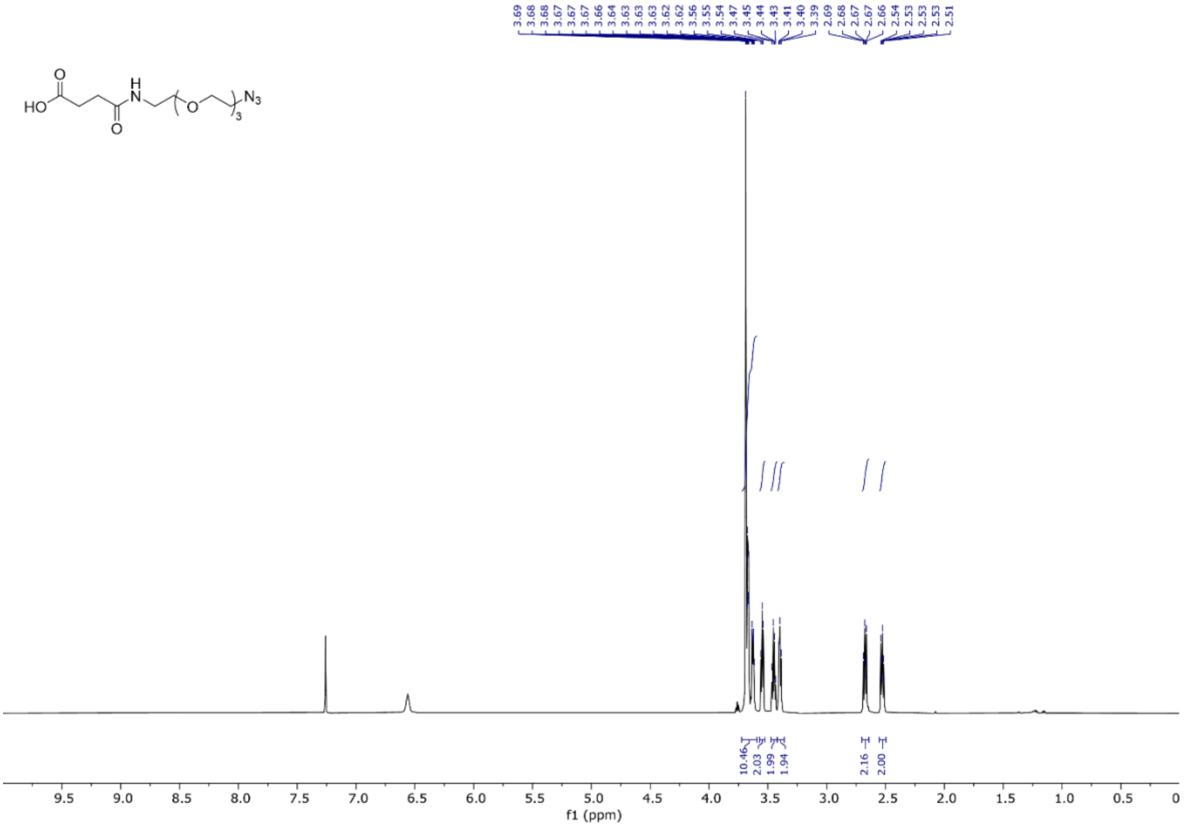


**Figure S18.** $^1$H NMR spectrum (400 MHz, $CDCl_3$), compound **S5**.

**Compound 1:**

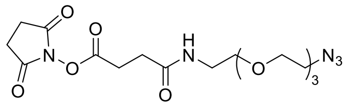


To a stirred solution of **S5** (467 mg, 1.47 mmol) in anhydrous DCM (6 mL), NHS (253 mg, 2.20 mmol) and EDC·HCl (422 mg, 2.20 mmol) were added at room temperature. The reaction was stirred for 2 h at room temperature, then quenched with $H_2O$ (30 mL) and extracted with DCM (5 x 20 mL). The combined organic layers were washed with brine (20 mL), dried with anhydrous $MgSO_4$, filtered and concentrated under reduced pressure. The residue was purified by column chromatography on silica gel ($CHCl_3$/MeOH 1:0 to 95:5, v/v) furnishing **1** (564 mg, 93 % yield) as a transparent syrup. $^1$H NMR (500 MHz, Chloroform-*d*) δ 3.70 – 3.60 (m, 10H. $OCH_2$), 3.58 – 3.53 (m, 2H, $OCH_2CH_2NH$), 3.46 (q, *J* = 5.2 Hz, 2H, $OCH_2CH_2NH$), 3.39 (t, *J* = 5.1 Hz, 2H, $N_3CH_2$), 2.99 (t, *J* = 7.3 Hz, 2H, $CH_2CH_2$), 2.83 (s, 4H, $CH_2^{NHS}$), 2.60 (t, *J* = 7.3 Hz, 2H, $CH_2CH_2$). NMR data are in agreement to those reported in the literature.[4]

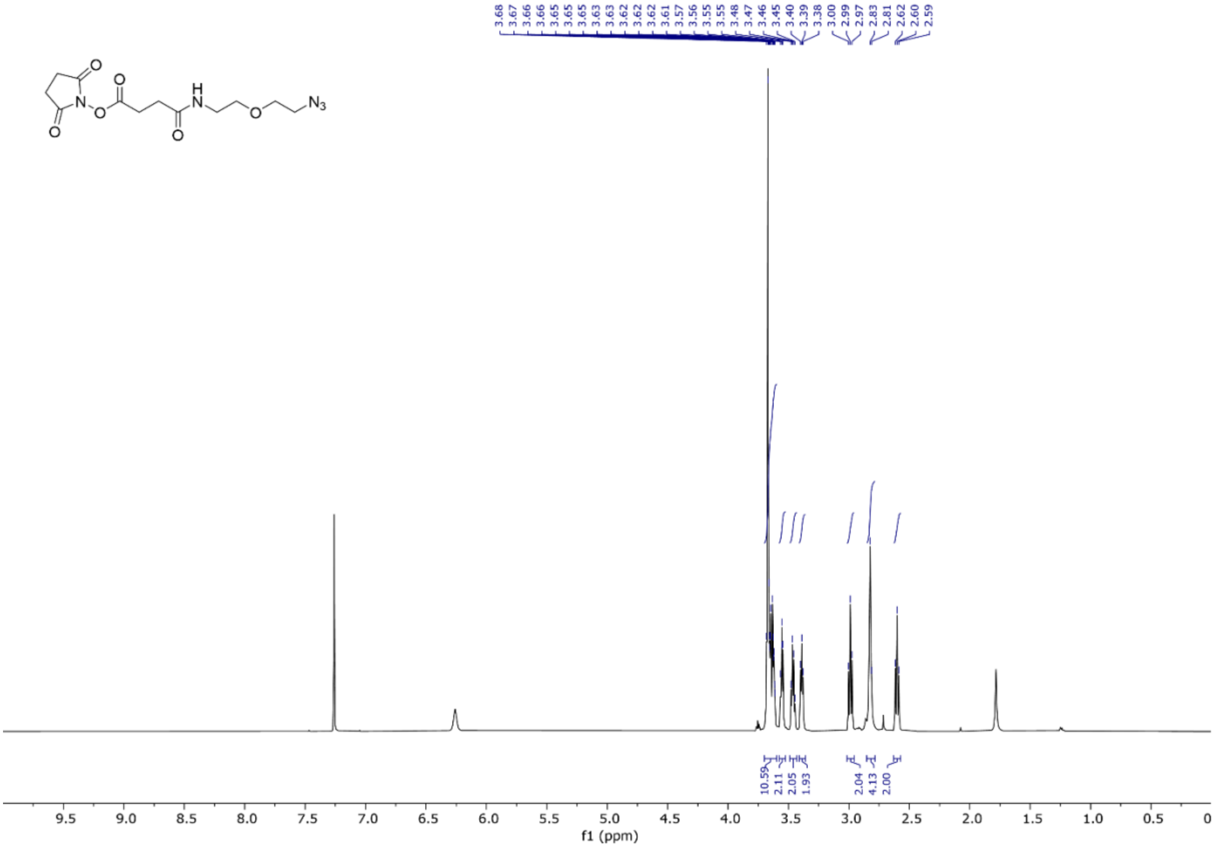

**Figure S19.** $^{1}H$ NMR spectrum (400 MHz, $CDCl_3$), compound **1**.

### 4. BSA and Abs functionalization

#### 4.1 preparation of BSA derivatives 10a-f

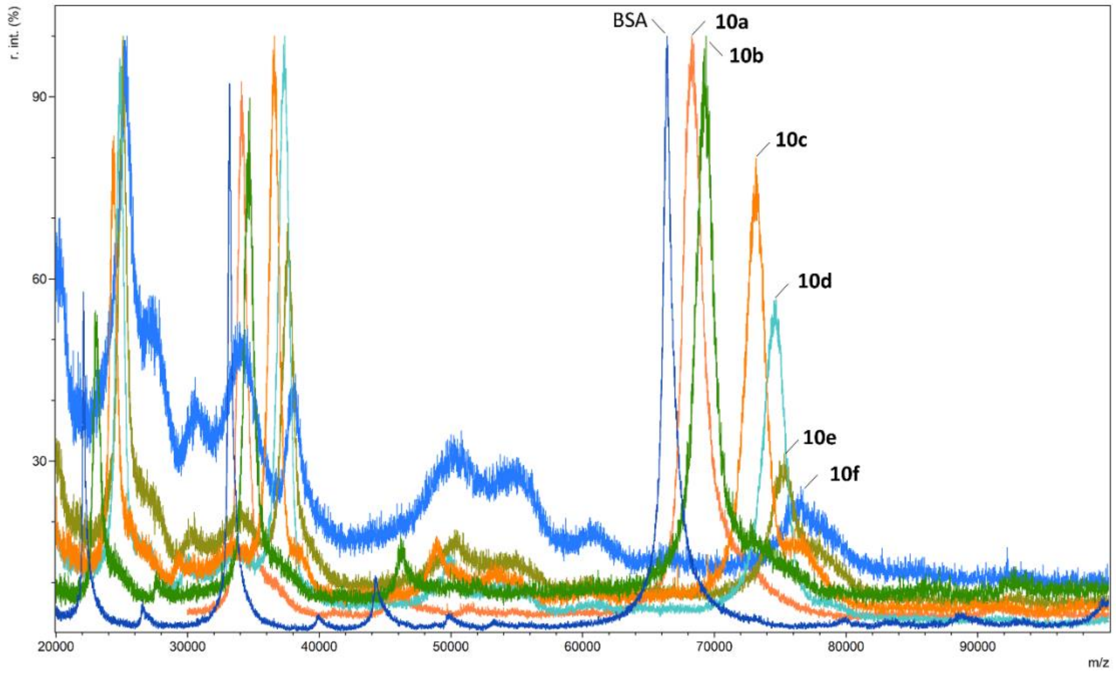


**Figure S20.** MALDI spectra of BSA and BSA-$N_3$ derivatives **10a-f**. Intensity is expressed as relative intensities (r. int.) %.

**Table S1.** Number of $N_3$ linkers incorporated on the BSA surface. Numbers were calculated based on the $M^{2+}$ peak.

| Compound | mg BSA | mmol BSA | mmol 1 | eq. of 1 | Approx. # of $N_3$ moieties |
|---|---|---|---|---|---|
| **10a** | 0.24 | $3.61 \cdot 10^{-6}$ | $0.14 \cdot 10^{-3}$ | 40 | 6.07 |
| **10b** | 0.24 | $3.61 \cdot 10^{-6}$ | $0.24 \cdot 10^{-3}$ | 67 | 10.36 |
| **10c** | 0.24 | $3.61 \cdot 10^{-6}$ | $1.20 \cdot 10^{-3}$ | 333 | 22.32 |
| **10d** | 0.24 | $3.61 \cdot 10^{-6}$ | $2.41 \cdot 10^{-3}$ | 666 | 27.26 |
| **10e** | 0.24 | $3.61 \cdot 10^{-6}$ | $3.61 \cdot 10^{-3}$ | 999 | 29.21 |
| **10f** | 0.24 | $3.61 \cdot 10^{-6}$ | $4.81 \cdot 10^{-3}$ | 1333 | 31.93 |

It is noteworthy that despite starting with a strong excess of linker (~ 40 eq.) only 10 linker units were incorporated on the protein surface. This may be related to the competing hydrolysis reaction of the NHS-activated esters in the slightly basic PBS buffer (pH 7.4). As expected, with a stronger excess of linker **1** more $N_3$ moieties were incorporated reaching a plateau of about 30 linker units. This indicates that the majority of the most solvent exposed primary amines reacted with **1**. The molecular peaks in the MALDI spectra showed a peak broadening proportional to the amount of **1** incorporated.

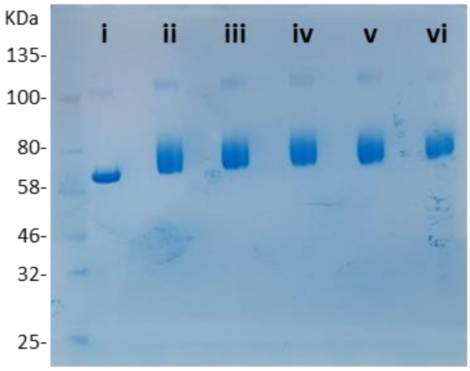

**Figure S22**. NuPAGE Gel electrophoresis of BSA derivatives **10b-f**: i) native BSA, ii) **10b**, iii) **10c**, iv)**10d**, v) **10e** and vi) **10f**.

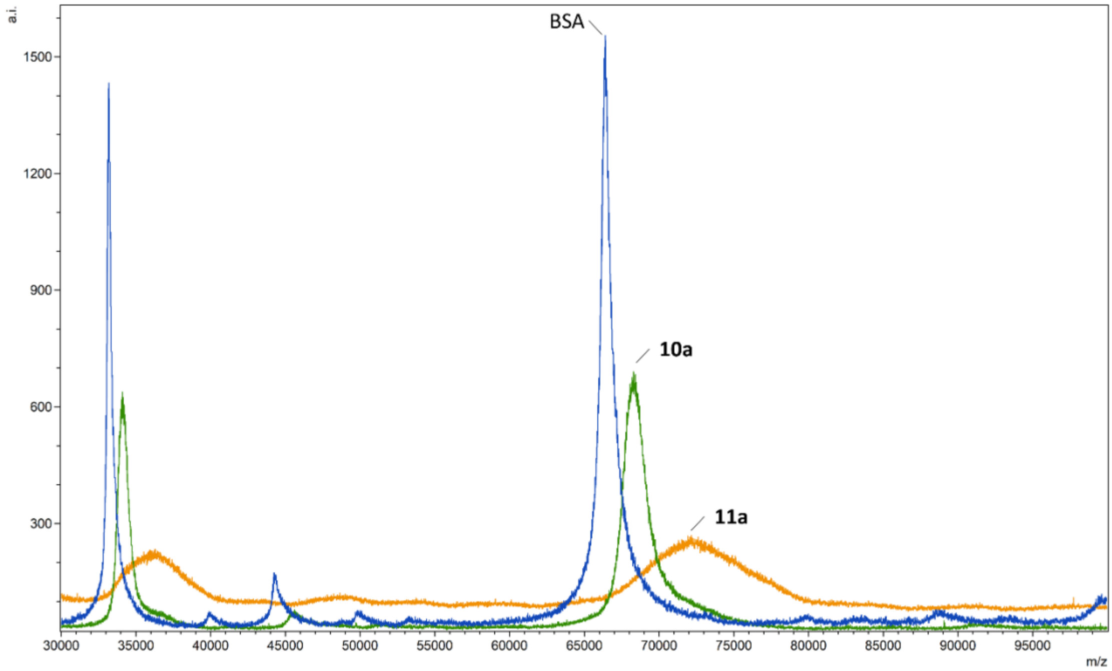

**Figure S23.** Full scale MALDI spectra of BSA, **10a**, and **11a**. Intensity is expressed as arbitrary units (a.i.).

#### 4.2. Preparation of Abs derivatives 12a-d and CD-Abs 13a-d

**Table S2.** Number of $N_3$ linkers incorporated on the Abs surface. Numbers were calculated based on the $M^{2+}$ peak.

| Compound | mg Abs | mmol Abs | mmol 1 | eq. of 1 | Approx. # of $N_3$ moieties |
|---|---|---|---|---|---|
| **12a** | 0.16 | $1.11{\cdot}10^{-6}$ | $0.12{\cdot}10^{-3}$ | 109 | 4.41 |
| **12b** | 0.16 | $1.11{\cdot}10^{-6}$ | $0.24{\cdot}10^{-3}$ | 217 | 8.57 |
| **12c** | 0.16 | $1.11{\cdot}10^{-6}$ | $1.20{\cdot}10^{-3}$ | 1086 | 26.36 |
| **12d** | 0.16 | $1.11{\cdot}10^{-6}$ | $2.41{\cdot}10^{-3}$ | 2172 | 30.21 |

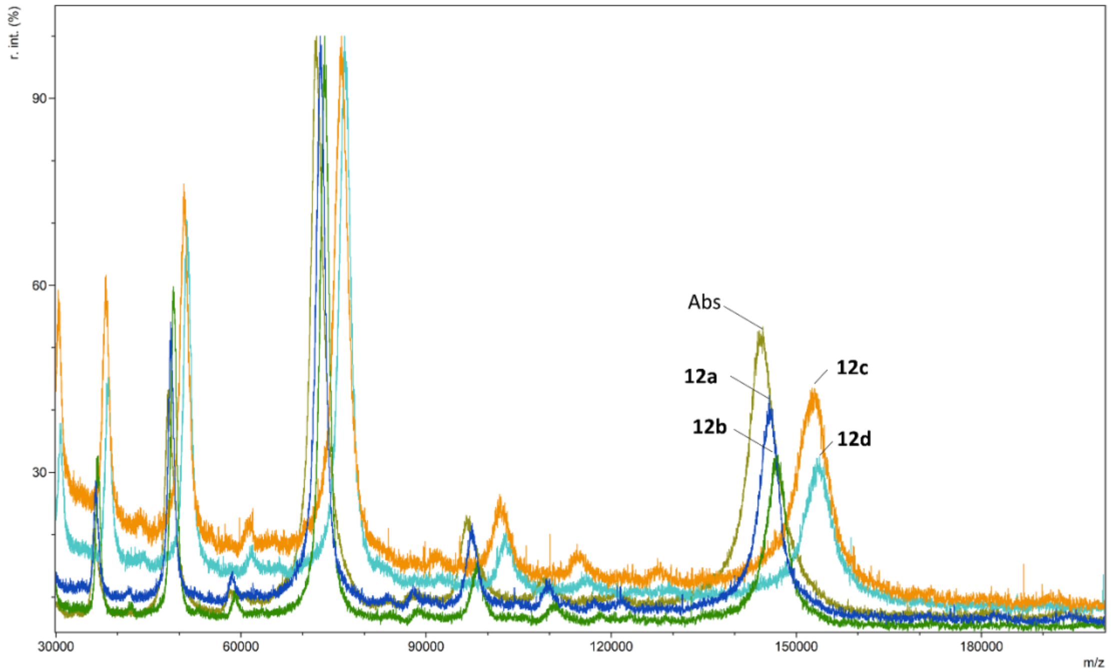


**Figure S24**. MALDI spectra of Abs and Abs-$N_3$ derivatives **12a-d**. Intensity is expressed as relative intensities (r. int.) %.

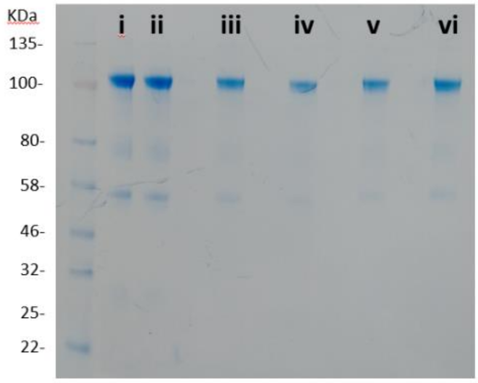


**Figure S25**. NuPAGE Gel electrophoresis of Abs derivatives **12a-d**: i) native Abs, ii) native Abs (no $N_3$ present) mixed with **2**, iii) **12a**, iv) **12b**, v)**12c** and vi) **12d**.

### 5. Microscopy images

#### 5.1. Optical microscope images

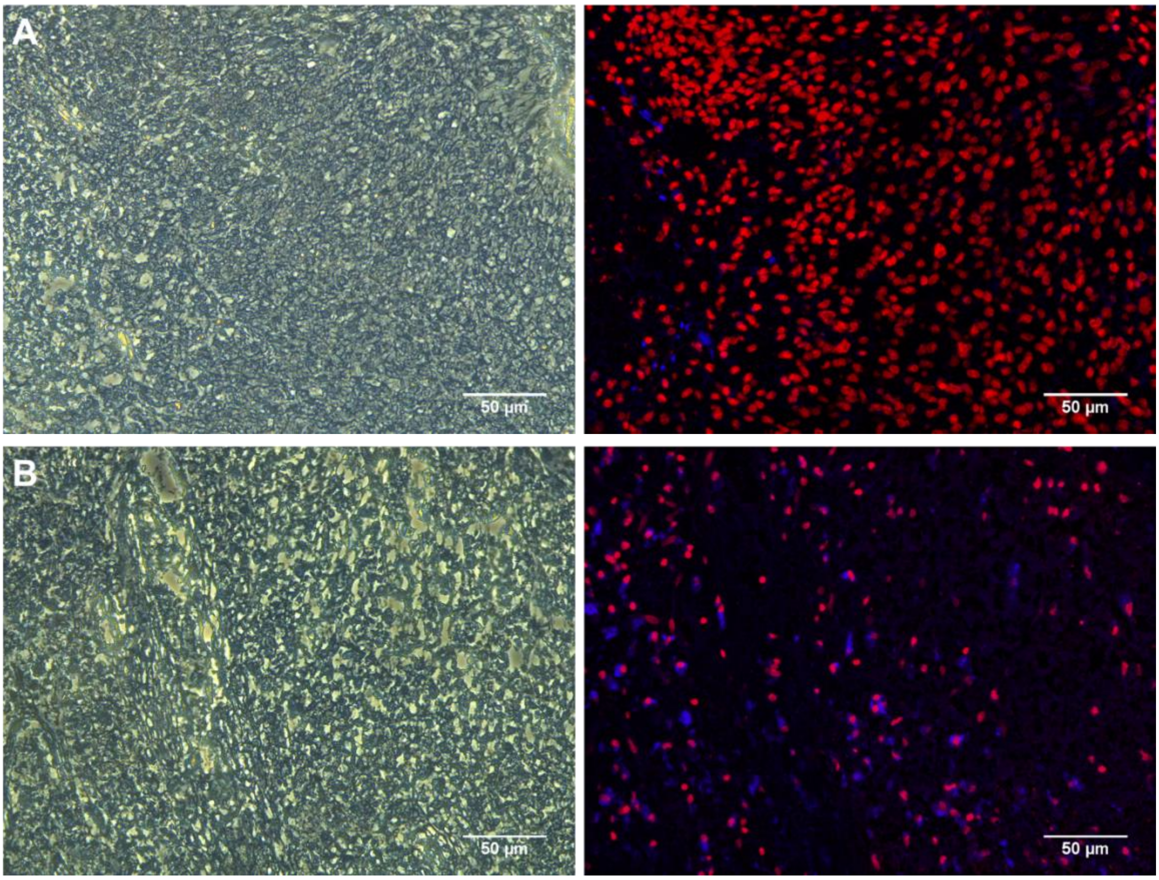


**Figure S26.** Optical microscope images of human tumour brain tissue. A) Negative control: tissue treated with **2** alone. B) Positive control: tissue treated with **13a**. Left: bright field and right: fluorescent channel. Nuclei are shown in red and immunostained regions are shown in blue.

#### 5.2. Confocal microscope images

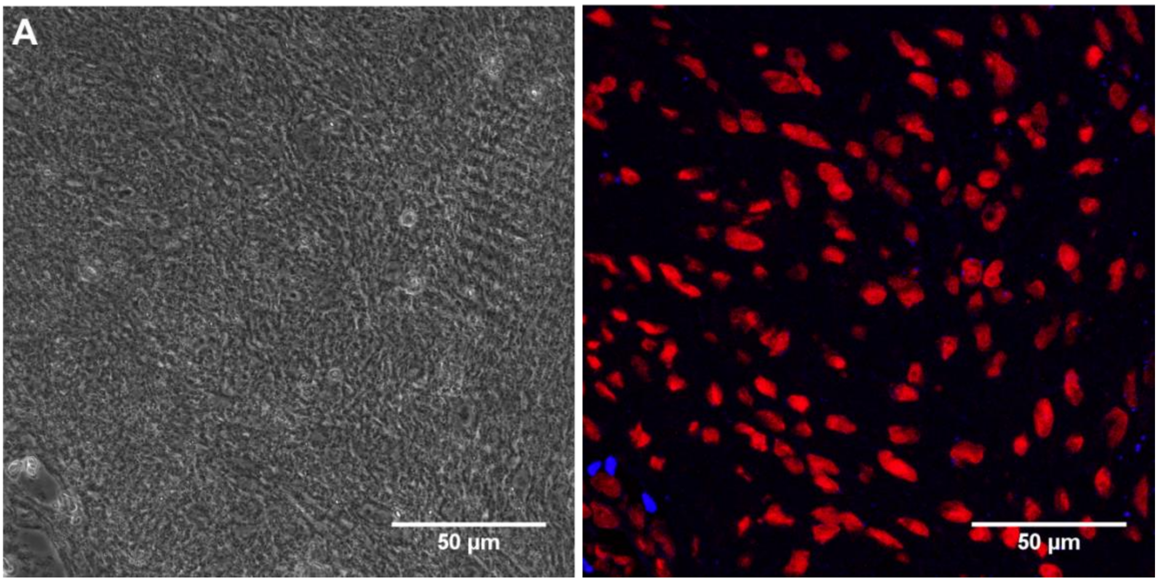

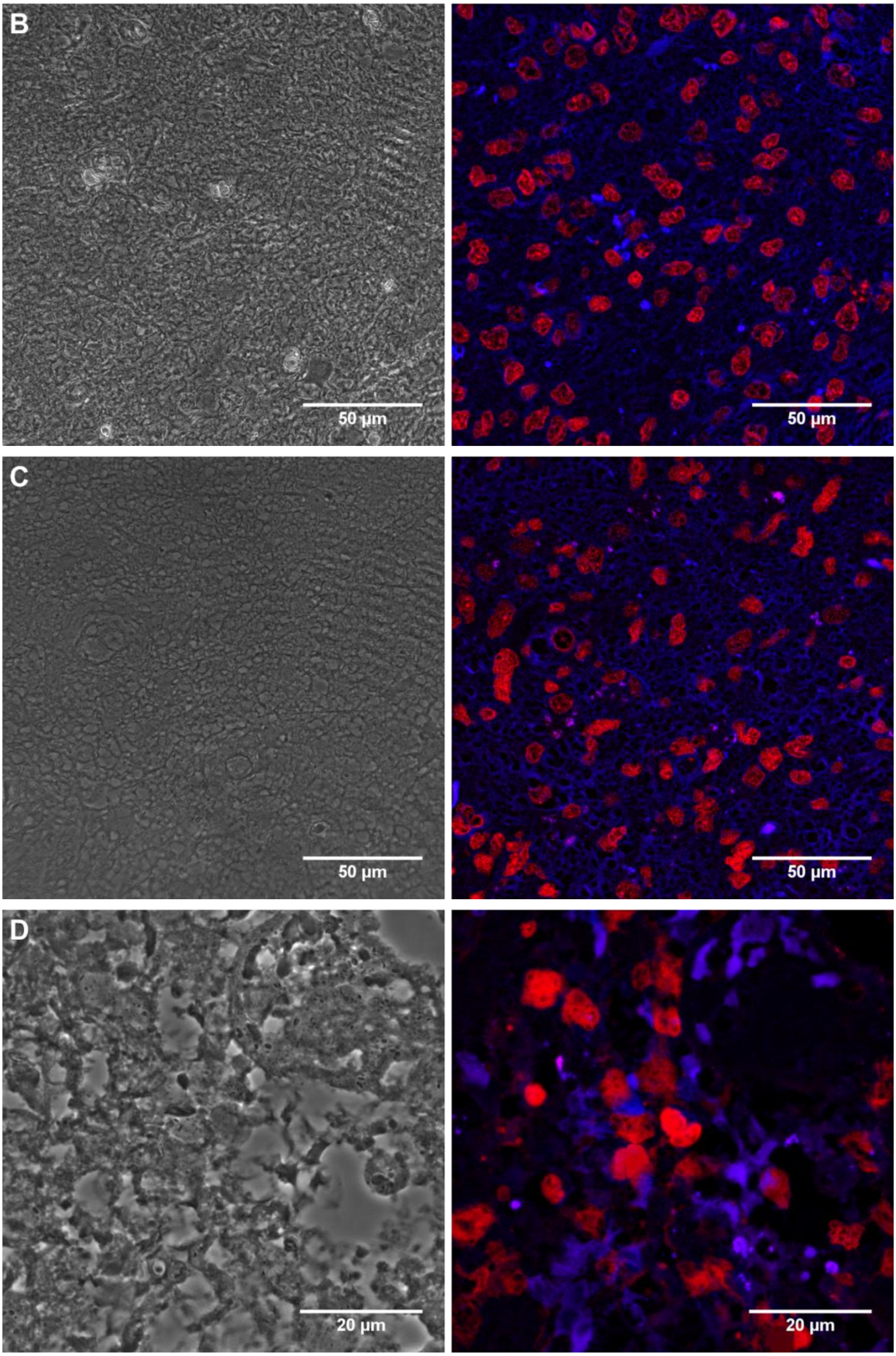
B
50 µm
50 µm
C
50 µm
50 µm
D
20 µm
20 µm

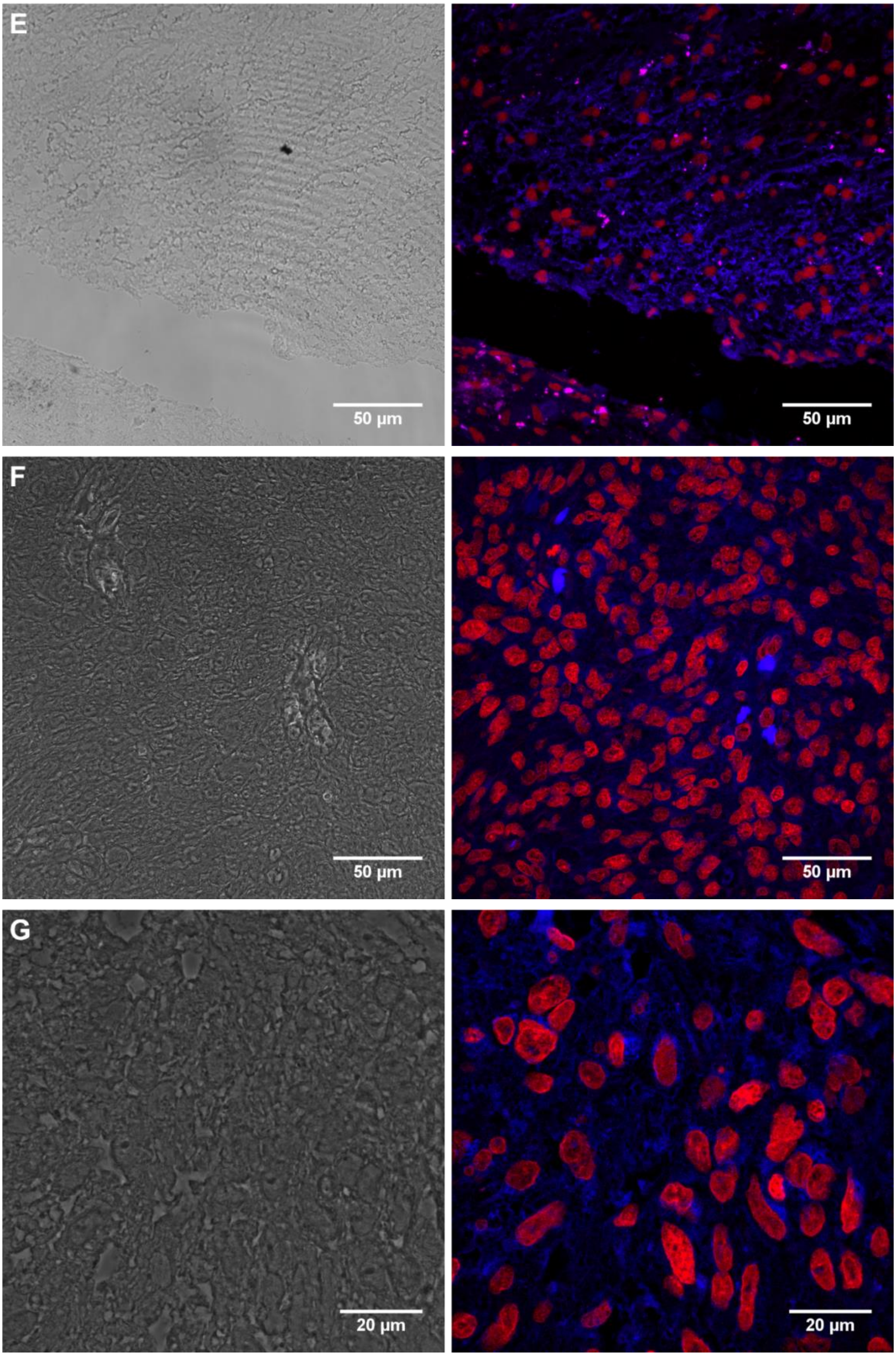
E
50 µm
50 µm
F
50 µm
50 µm
G
20 µm
20 µm

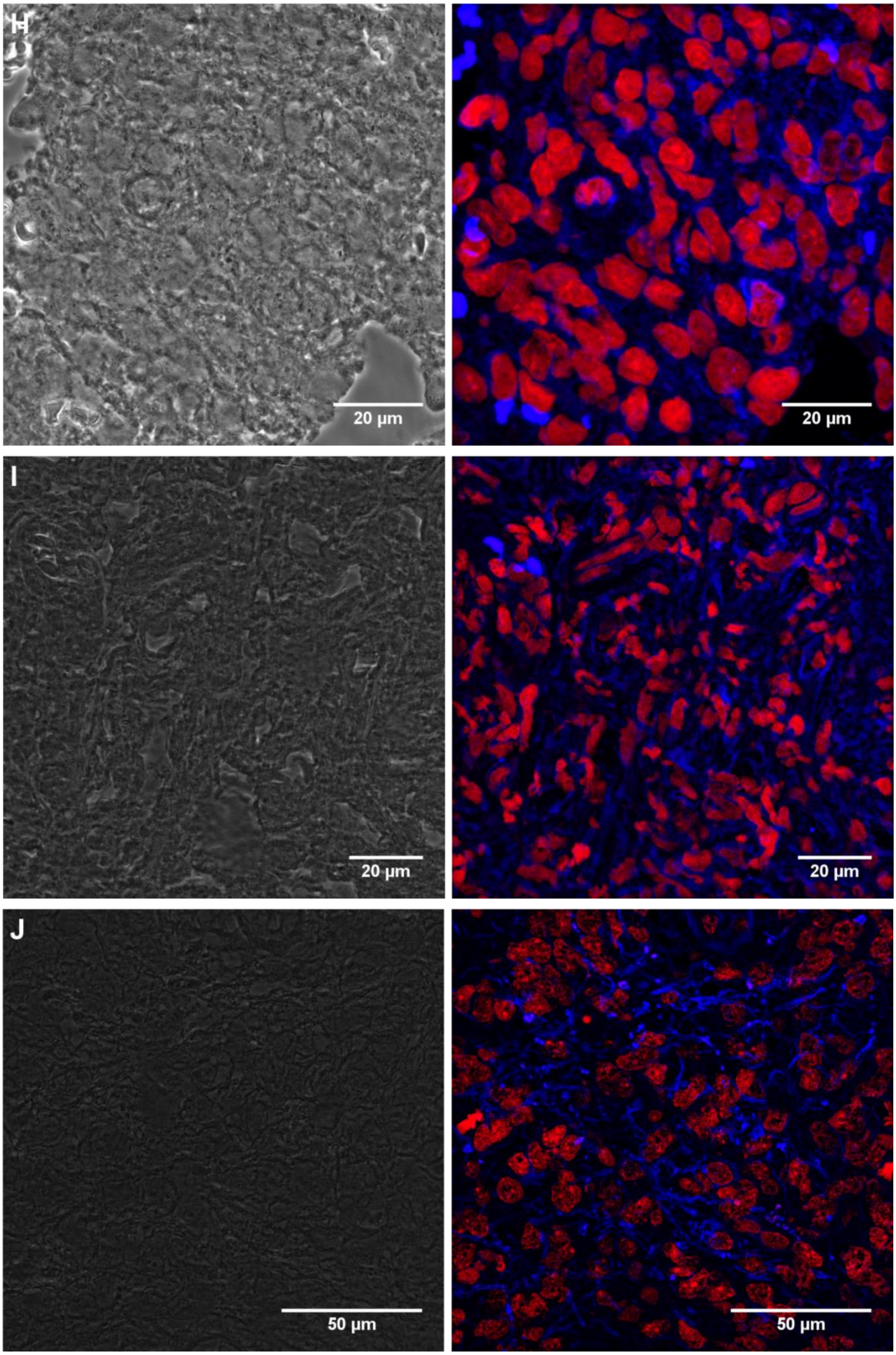
H
20 µm
20 µm
I
20 µm
20 µm
J
50 µm
50 µm

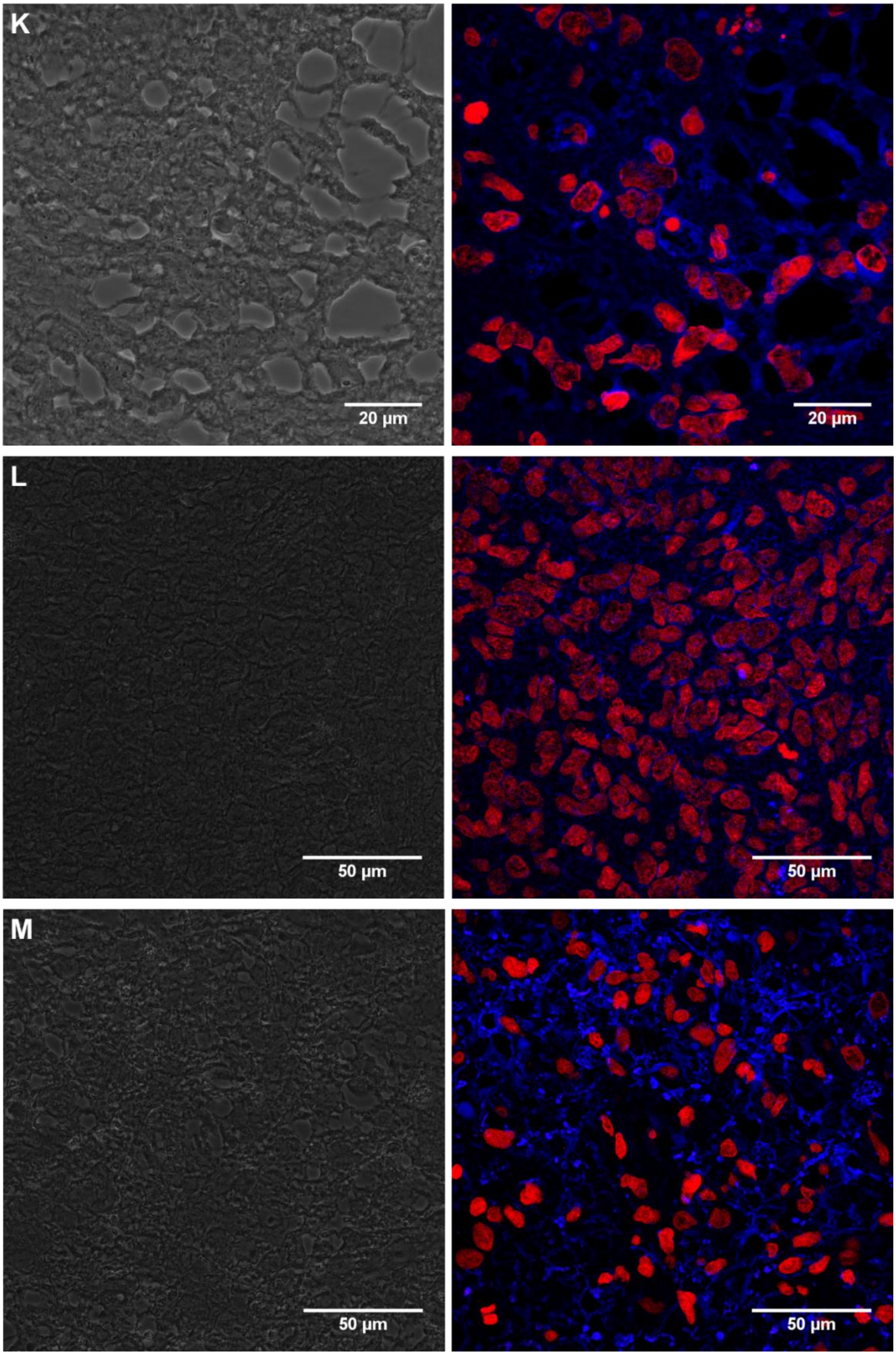


**Figure S27.** Optical microscope images of human tissues. A) Negative control: spinal tissue diagnosed as schwannoma (no GFAP expression) treated with **13a**. B-M) Positive controls: brain tumour tissues diagnosed as GBM treated with **13a**. Left panels: bright field showing monolayer and right panels: fluorescent channel of anti-GFAP Abs-CD probe **13a** labelled tissue. Nuclei are shown in red and immunostained regions are shown in blue.

#### 5.3. Clinical data for brain tumour samples

**Table S3**. Clinical data for for brain tumour tissues used for the immunofluorecence detection of GFAP.

| Image | Patient ID | Age | Sex | Site | Diagnosis | WHO grade | IDH status | Legacy # |
|---|---|---|---|---|---|---|---|---|
| **S26 A** | 13/1017A | 55 | M | left frontal | GBM | 4 | negative(i) | 13N90033869 |
| **S26 B** | 13/1017A | 55 | M | left frontal | GBM | 4 | negative(i) | 13N90033869 |
| **S27 A** | 21/N1214A1 | 76 | M | spinal L5/s1 | schwannoma | 1 | n.s. | n.s. |
| **S27 B** | 13/1017A | 55 | M | left frontal | GBM | 4 | negative(i) | 13N90033869 |
| **S27 C** | 07/1283B | 64 | M | left frontal | GBM | 4 | | 07N90026012 |
| **S27 D** | 07/0257 | 65 | M | right parieto-occipital | GBM | 4 | n.s. | 07N90025010 |
| **S27 E** | 12/0141 | 63 | F | left parietal | GBM | 4 | n.s. | 12N90026444 |
| **S27 F** | 08/0054B | 70 | M | right parietal | GBM | 4 | n.s. | 08N90026180 |
| **S27 G** | 10/0053* | 52 | M | right parietal | GBM | 4 | negative(i,s) | 10N90029111 |
| **S27 H** | 11/0111A | 68 | M | right temporal | GBM | 4 | negative(i) | 11N90030504 |
| **S27 I** | 08/0037 | 67 | F | right frontal | GBM | 4 | n.s. | 08N90026160 |
| **S27 J** | 08/0057B | 58 | M | n.s. | GBM | 4 | n.s. | 08N90026183 |
| **S27 K** | 08/0160B | 55 | M | right hemisphere | GBM | 4 | n.s. | 08N90026275 |
| **S27 L** | 10/0452 | 42 | F | multifocal | GBM | 4 | n.s. | 10N90029421 |
| **S27 M** | 10/0865 | 70 | M | right temporal | GBM | 4 | n.s. | 10N90029881 |

n.s.: not stated. Entries are related to microscope images, section 5.1 and 5.2. * additional available infomration for patient 10/0053: MGMT status: methylated; BRAF V600E: negative; TERT: mutated c.-124C>T.
